# Interpersonal heart rate synchrony predicts effective information processing in a naturalistic group decision-making task

**DOI:** 10.1101/2023.07.24.550277

**Authors:** K. M. Sharika, Swarag Thaikkandi, Nivedita, Michael L. Platt

## Abstract

Groups often outperform individuals in problem-solving. Nevertheless, failure to critically evaluate ideas risks sub-optimal outcomes through so-called *groupthink*. Prior studies have shown that people who hold shared goals, perspectives or understanding of the environment show similar patterns of brain activity, which itself can be enhanced by consensus building discussions. Whether shared arousal alone can predict collective decision-making outcomes, however, remains unknown. To address this gap, we computed interpersonal heart rate synchrony, a peripheral index of shared arousal associated with joint attention, empathic accuracy and group cohesion, in 44 groups (n=204) performing a collective decision-making task. The task required critical examination of all available information to override inferior, default options and make the right choice. Using multi-dimensional recurrence quantification analysis (MdRQA) and machine learning, we found that heart rate synchrony predicted the probability of groups reaching the correct consensus decision with greater than 70% cross-validation accuracy—significantly higher than that predicted by the duration of discussions, subjective assessment of team function or baseline heart rates alone. We propose that heart rate synchrony during group discussion provides a biomarker of interpersonal engagement that facilitates adaptive learning and effective information sharing during collective decision-making.

## 1 Introduction

Findings across multiple research domains, including organizational psychology (Woolley *et al.,* 2010) and ecology (Sasaki *et al.,* 2013), have highlighted the potential of groups to outperform individuals in solving complex problems. This collective advantage is thought to reflect the pooling of diverse skills and experiences of group members to resolve a given challenge. However, failure to critically re-examine ideas may lead to sub-optimal outcomes, as in *groupthink* wherein individual members’ “strivings for unanimity override their motivation to realistically appraise alternative courses of action” (Janis, 1972;Mayton & Zachary Brink, 2011). Multiple studies have identified similarity in brain activity between individuals as a biomarker of shared goals, perspectives or understanding of the environment (Cui *et al.,* 2012; Lahnakoski *et al.,* 2014; Golland *et al.,* 2015; Yeshurun *et al.,* 2017; Hu *et al.,* 2018; Nguyen *et al.,* 2019). Collective task performance is predicted by such interbrain synchrony (Reinero *et al.,* 2021) which can be enhanced by consensus building discussions (Sievers *et al*.). Interpersonal heart rate synchrony, a peripheral index of shared arousal, has been associated with joint attention, empathic accuracy and group cohesion (Levenson & Gottman; 1983; Levenson & Ruef, 1992; Konvalinka *et al.,* 2011; Mitkidis *et al.,* 2015; Jospe *et al.,* 2020; Gordon *et al.,* 2021; Tomashin *et al.,* 2022). Whether heart rate synchrony between group members can predict the quality of group discussion outcomes remains unknown (Haynes & Platt, 2022; Aldag & Fuller; 1993).

To address these questions, we measured heart rates in three-to-six member groups performing a collective decision-making task (Fig. 1) based on the Hidden Profile Paradigm (Stasser & Titus, 1985; Larson Jr 1997; De Wilde *et al.,* 2017) in which information necessary for solving a problem is distributed unequally among group members. Specifically, participants faced the challenge of selecting the best person for a university faculty position based on attributes provided to them about three potential candidates. While the problem had an objective correct answer, it was designed to favor one of the incorrect options if individuals were to either go by the information provided to them alone or that was shared between all group members (Table 1). In other words, successful performance in the task required active consideration and critical examination of all available information between members to override the inferior, default options. By collecting and analyzing heart rate data of groups while they performed this task, we tested if interpersonal heart-rate synchrony could predict the effectiveness of groups in overriding the inferior option. We hypothesized that since critical evaluation of unique information would entail expression (and consequent resolution) of differences of opinion among group members, heart rate synchrony would distinguish between groups that promoted such interpersonal risk-taking among members and those that did not. The tendency for interpersonal risk-taking in teams, conceptualized as team psychological safety, is known to be an essential component of adaptive learning in groups (A. Edmondson, 1999). Since psychological safety is likely facilitated via empathic listening and perspective-taking (Kluger & Itzchakov, 2022), processes that have been previously associated with high interpersonal physiological synchrony (Di Mascio *et al.,* 1955; Stratford *et al.,* 2012; Chatel-Goldman *et al.,* 2014; Marci, 2006; Ritov & Richter-Levin, 2014; Messina *et al.,* 2013, Jospe *et al.,* 2020; Levenson & Ruef, 1992), we predicted that successful performance in this task would be associated with enhanced heart rate synchrony between group members.

**Figure 1:**
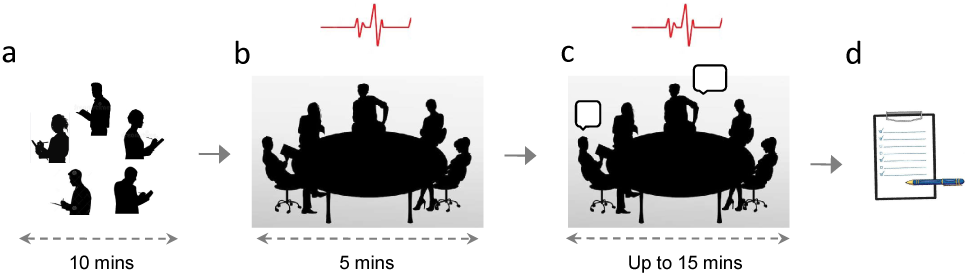
Schematic of Experimental Procedure. The group decision-making task simulated choosing the best candidate for a new faculty position at a university. a) Participants were provided 10 minutes to first read the information sheet summarizing the attributes of three potential candidates on their own, while making written notes to support ensuing discussion. Heart rate data was collected while participants were b) sitting and not talking (preGD) for 5 minutes c) involved in the group discussion (GD) which lasted for a maximum of 15 minutes across groups. d) Following the discussion, each participant had to individually respond to a set of questionnaires (GQ) on their experience of working with the group. Image used is adapted from www.freepik.com.

**Table 1:** Outline of the structure of the task by showing how the information units (IUs, 24 in total) related to the three candidates, A, B and C (associated with 6, 9, 9 IUs, respectively), were distributed across group members (M1 to M6). While IUs in black (A1-A3 about candidate A, B1-B6 about candidate B and C1-C6 about candidate C) are positive attributes shared with all members of the group, IUs in green and red are positive and negative attributes, respectively, provided only to select members (M1 to M3). IUs are distributed such that group members are likely to choose candidate B or C as their preferred candidate, based on data provided to them. However, candidates B and C, with overall 6 positive and 3 negative attributes each, are inferior choices to this best candidate selection problem. Only when unique data provided to group members M1 to M3 are introduced and critically considered in the discussion, would the correct answer (candidate A with no negative attributes) emerge.

| M1 | M2 | M3 | M4 | M5 | M6 |
| --- | --- | --- | --- | --- | --- |
| A1,<br>A2,<br>A3,<br>A4 | A1,<br>A2,<br>A3,<br>A5 | A1,<br>A2,<br>A3,<br>A6 | A1,<br>A2,<br>A3 | A1,<br>A2,<br>A3 | A1,<br>A2,<br>A3 |
| B1,<br>B2,<br>B3,<br>B4,<br>B5,<br>B6,<br>B7 | B1,<br>B2,<br>B3,<br>B4,<br>B5,<br>B6,<br>B8 | B1,<br>B2,<br>B3,<br>B4,<br>B5,<br>B6,<br>B9 | B1,<br>B2,<br>B3,<br>B4,<br>B5,<br>B6 | B1,<br>B2,<br>B3,<br>B4,<br>B5,<br>B6 | B1,<br>B2,<br>B3,<br>B4,<br>B5,<br>B6 |
| C1,<br>C2,<br>C3,<br>C4,<br>C5,<br>C6,<br>C7 | C1,<br>C2,<br>C3,<br>C4,<br>C5,<br>C6,<br>C8 | C1,<br>C2,<br>C3,<br>C4,<br>C5,<br>C6,<br>C9 | C1,<br>C2,<br>C3,<br>C4,<br>C5,<br>C6 | C1,<br>C2,<br>C3,<br>C4,<br>C5,<br>C6 | C1,<br>C2,<br>C3,<br>C4,<br>C5,<br>C6 |

## 2 Methods

Discrete heart rate data (beats per minute for every 2 seconds) was collected from 271 participants (88 males, 182 females) spanning the ages 18-35 years (mean ± SEM = 22 ± 0.22) using wireless sensors (Polar H10 chest straps). Participants were grouped into 58 teams of three to six members each (9 groups with n=3, 15 groups n=4, 20 groups with n=5, 14 groups with n=6) with no two members in the same group self-reporting prior familiarity with each other. The heart rate sensor’s proprietary application, Polar Club, permitted simultaneous recording from all members of each team for both baseline (sitting and not talking before the group discussion, preGD data, for 5 mins) and task-related (group discussion, GD data, for a maximum of 15 mins) epochs. All participants gave informed consent to participate in the experiment. The study was approved by the Institutional Review Board of the University of Pennsylvania, Philadelphia, USA.

### Task

The group decision-making task (Fig. 1) simulated choosing the best candidate for a new faculty position at a university. Participants were provided 10 minutes to first read the information sheet summarizing the attributes of three potential candidates on their own, while making written notes to support ensuing discussion (but not for direct perusal of other group members). They were explicitly told that only one candidate was most suitable for the position and that when considering different attributes between candidates, they should value each attribute equally, even though these attributes might be qualitatively different from each other. Unknown to participants, information about the candidates was distributed unequally among group members. Some information units were presented only to some team members (unique information units) while other information units were distributed to all team members (shared information units).

Table 2 describes the distribution of information across team members. Since unique information was crucial for arriving at the correct answer, correct consensus decisions required all team members to introduce all the information available to them into discussion, and to critically evaluate each candidate’s positive and negative attributes, before making a final selection. Groups were permitted a maximum of 15 minutes to discuss and render a decision, while their deliberations were recorded for both audio and video. After group discussion, each participant answered the following questionnaires on their experience of working with their group: 1) the Edmondson 7-item survey on group psychological safety (A. Edmondson, 1999; A. C. Edmondson *et al.,* 2004); 2) the three scales of the Team Diagnostic survey (Wageman *et al.,* 2005): a) process criteria for team effectiveness b) team interpersonal processes and 3) individual learning and well-being.

**Table 2:** Description of RQA variables computed from the Recurrence Plot.

| RQA variables | Description |
| --- | --- |
| Recurrence Rate (RR) | Measures density of recurrence points in the recurrence plot, indicating how probable recurrence of states is in the system. |
| Determinism (DET) | Measures what fraction of the diagonal line lengths are above a minimum, given, line length. $p(l)$ is the probability of a line length $l$. Since diagonal lines are markers of consecutive periods of recurrence in the data, determinism corresponds to the predictability of the dynamical system. |
| Laminarity (LAM) | Mathematically equivalent to determinism but defined for vertical (or horizontal) line lengths. Since vertical (or horizontal) lines are markers of states that do not change or change very slowly, laminarity quantifies the extent of the dynamical system being trapped in any given state for some time (Marwan, Romano, *et al.*, 2007). |
| Average diagonal line length (L) | Average value of diagonal line length distribution, quantifying how far in time the dynamical system is predictable. |
| Average vertical line length/ Trapping time (TT) | Average value of the vertical line length distribution. |
| Maximum diagonal line length | Maximum value from the diagonal line distribution |
| Maximum vertical line length | Maximum value from the vertical line distribution |
| Diagonal and Vertical Entropy (ENTR) | Quantifies the degree of uncertainty in the possible states and hence, the complexity of the dynamical system, using the distribution of diagonal and vertical line lengths present in the plot, respectively. |

### Data Analysis

#### Multi-dimensional Recurrence Quantification Analysis (MdRQA)

Recurrent quantification analysis (RQA) quantifies the pattern of recurrence (or how often a particular value is repeated over time) in the phase space of a dynamic system (Webber Jr & Zbilut, 1994; Marwan, Wessel, *et al.,* 2002). A phase space is a collection of all possible states of the system plotted as a function of time. How often a trajectory revisits a point in the phase space, or recurrence, indicates how different components of a multi-variate system interact and converge on the same state across time (Mitkidis *et al.,* 2015). The phase space of a multidimensional system can be reconstructed using Taken’s time delayed embedding theorem (Takens, 1981), which states that if one has access to only one variable from a complex system governed by multiple interdependent variables, then one can reconstruct the dynamics of this system by utilizing (D number of) time delayed versions of the observable x as (D-dimensional) coordinates of the phase space (Huffaker *et al.,* 2017). Let the time series be:

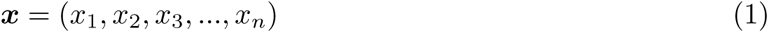

which is sampled in regular intervals in time. From this, D-dimensional vectors, ***V*_1_**, ***V*_2_**, and so on, can be constituted by estimating D-1 versions of ***x*** with time delay, *τ* , as below:

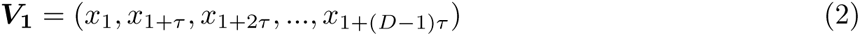

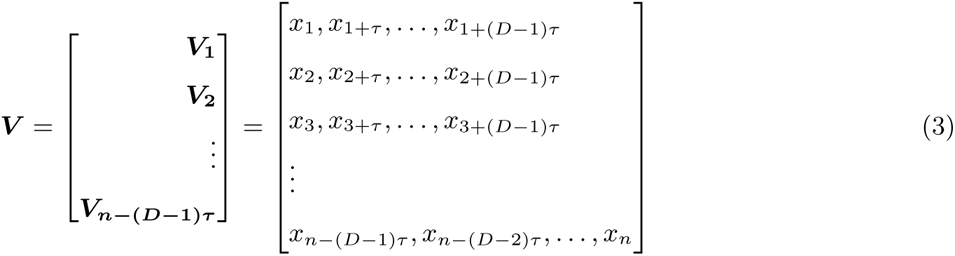

In the above expression, while each row represents a point in the D-dimensional phase space, each column represents the evolution of the trajectory in one of the D dimensions. MdRQA is a multidimensional extension of RQA which takes the time series of each of the group members as independent dimensions and quantifies recurrence at the group level (Wallot, Roepstorff, *et al.,* 2016; Gordon *et al.,* 2021; Tomashin *et al.,* 2022).

Heart rate data (beats per minute) was z-transformed for the MdRQA analysis (Gordon *et al.,* 2021) since we were interested in how it changed with respect to members in a group over time rather than the absolute values. To generate the recurrence plots, time delay & embedding dimension were computed for each group separately. Time delay (tau, *τ* ) was estimated by serially sampling values from 1 to 20 seconds and computing the first minima (first local minima or global minima when the local minima did not exist) of the mutual information between the time series of a group and a time delayed version of it (Bevilacqua *et al.,* 2019). This approach ensures that the time delayed signals are not too similar and permits the multidimensional topology of the trajectories in the phase space to unfold completely (see Kantz & Schreiber(2004), section 9.2, page 150 and supplementary section D for more details). For our data, tau ranged from 1 to 9 across groups. The number of embedding dimensions, m, required to adequately reconstruct the phase space was estimated using the false nearest neighbor approach (Kennel *et al.,* 1992; Hegger & Kantz, 1999; see Kantz & Schreiber(2004), section 3.3.1, page 37, figure 3.3). For our data, m ranged from 3 to 13 across groups. We chose the threshold radius, epsilon, which decides how close two points in the phase space need to be to be considered recurrent, to be such that the recurrence rate (percentage of recurrent points/ black dots in the recurrence plot) was constant across groups (10%). This allowed other variables derived from the distribution of black dots in the recurrence plot (percentage of diagonal lines (determinism) and percentage of vertical lines (laminarity), average diagonal & average vertical line, maximum diagonal & maximum vertical line, diagonal and vertical entropy; see Table 2 for their respective definitions) to be directly comparable across groups. For our data, epsilon ranged from 1.7 to 7.7 across groups. Figure 2 illustrates example recurrence plots for two groups: one that reached a correct consensus (left panel) and one that did not (right panel).

**Figure 2:**
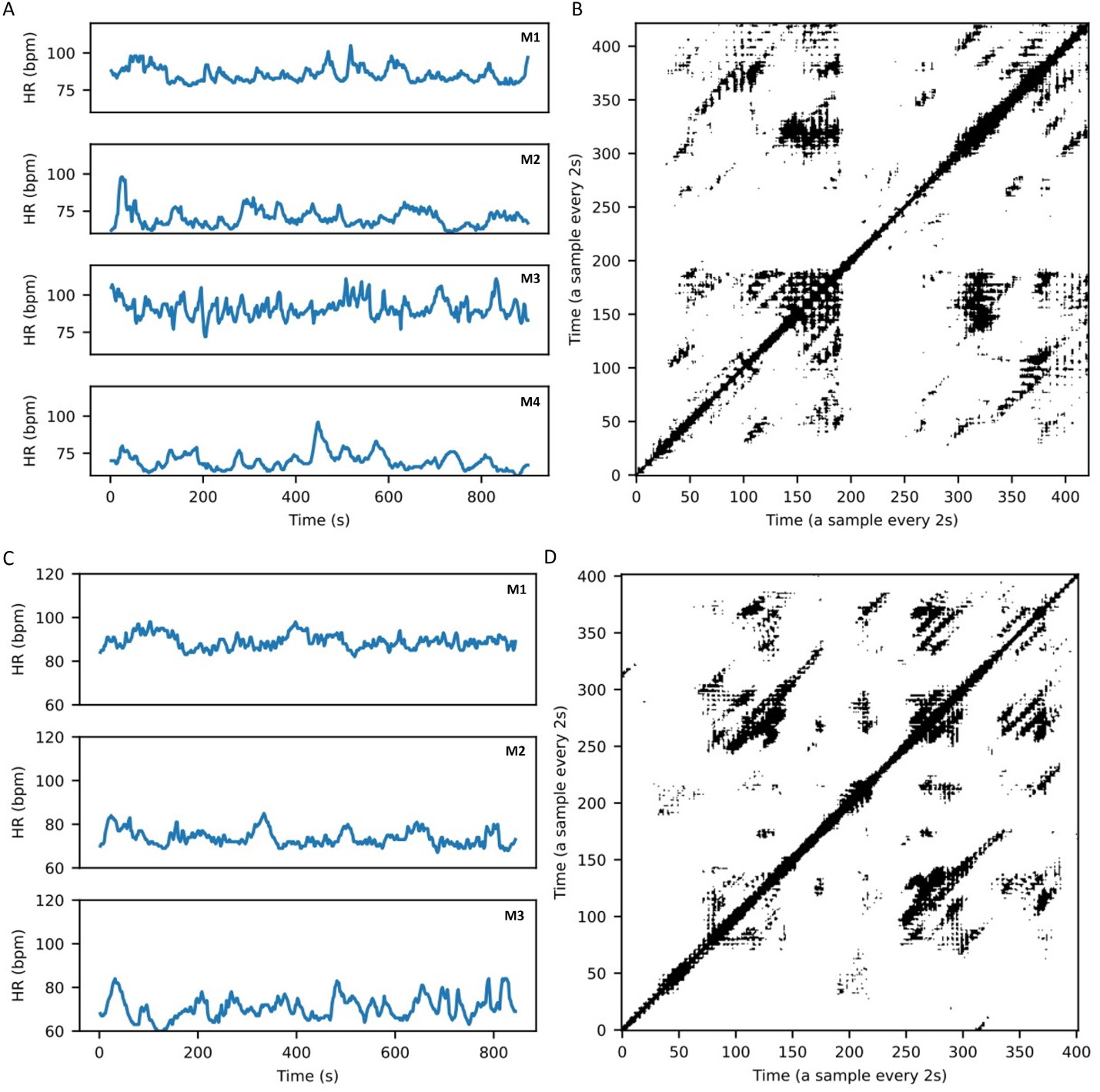
Raw heart rate data of individual members (M1-M4 in A and M1-M3 in C) and recurrence plot (B and D) for an example group that did not arrive at the correct consensus outcome (top panel) and for one that arrived at the correct consensus outcome (bottom panel). HR is collected at 0.5Hz

**Figure 3:**
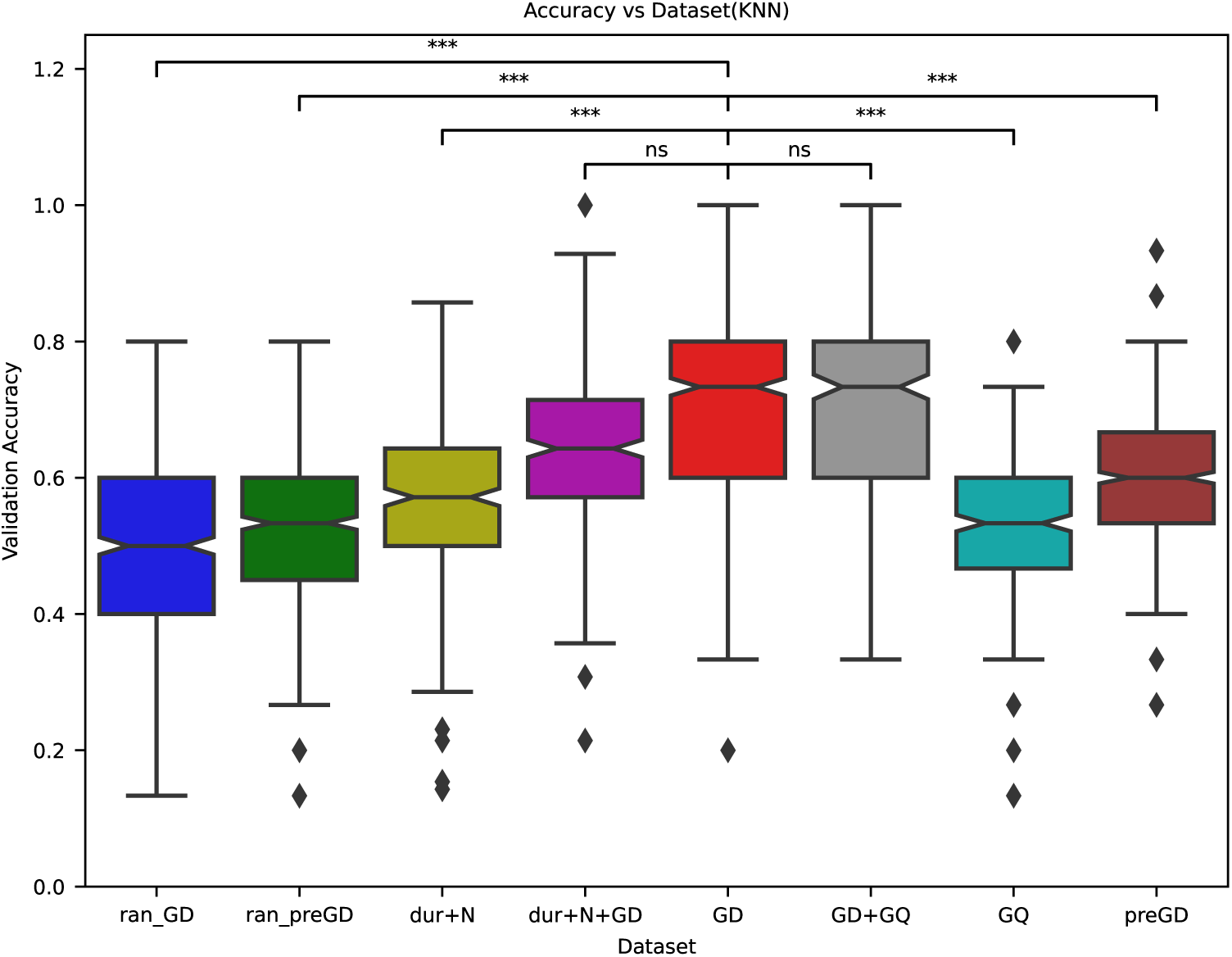
Classifier performance, represented by the cross-validation accuracy for discriminating between groups that reached a correct consensus vs. not, was compared when it was trained using the following datasets: (a) MdRQA measures from time series data during the group discussion (GD) (b) MdRQA measures from time series data during baseline (preGD) (c) MdRQA measures from randomized time series data during group discussion (ran-GD) (d) MdRQA measures from randomized time series data during baseline (ran-preGD) (e) average group response to a triad of questionnaires on subjective group experience post-discussion (GQ) (f) MdRQA measures from time series data during the group discussion as well as average group questionnaire response post-discussion (GD+GQ) (g) Duration (dur) and group size (N) data (dur+N) (h) MdRQA measures from time series data during the group discussion as well as duration and group size data (dur+N+GD). Identity of groups going into training and validation fold, respectively, was preserved across datasets for creating a distribution of accuracy for comparison using Wilcoxon signed-rank test.. MdRQA measures refer to the mode of the MdRQA variable distribution across sliding windows of a group RP

### Sliding window MdRQA followed by Nested Cross-validation of KNN classification

Since data collection was discontinued when groups reached a consensus, the durations of group discussion varied across groups in our data set (mean ± SEM: 8.6 mins ± 36s) and resulted in recurrence plots (RPs) of different lengths. Unequal sized RPs cannot be compared directly using linear scaling because individual RPs may be sensitive to local dynamics to different extents. For the same reason, while shorter RPs may capture local dynamics, the latter may get averaged out in longer RPs. Therefore, to compare recurrent plots of different sizes, we used a sliding window approach followed by nested cross-validation of a K-Nearest Neighbour (KNN) classification procedure (k=5) that has together been shown to classify binary states of a non-linear dynamical system across varying levels of noise with high cross-validation accuracy (Thaikkandi & Sharika, 2023). A non-parametric, distance-based method of classification like KNN was used instead of using a regression model because we were interested in broadly testing if and how MdRQA variables (that are often correlated with each other for discrete heart rate data (Gordon *et al.,* 2021) predict group decision outcomes rather than studying how each variable accounts for the variability in decision outcomes individually.

The sliding window approach involves partitioning a single large RP into smaller, same sized, overlapping windows along the RP diagonal and then computing the RQA variables for each of these sliding windows. Since RQA variables are statistical measures (e.g., average, max, entropy etc.) obtained from histogram distributions of vertical or horizontal lines of different lengths in the RP, it is important that the time window being examined for generating these histogram distributions be large enough to estimate the local dynamics with low variance and high statistical confidence. To select the time window that can be used to compare RPs of unequal sizes, we used a bootstrapping method (Marwan et al., 2013) with sliding windows of different lengths to first estimate the lowest sliding window size likely to give MdRQA variable estimates with high statistical confidence. We chose a common minimum sliding window size of 68 across all groups (see Supplementary section C for details) to first compute the MdRQA variables for each sliding window, resulting in a distribution of these variables across sliding windows for a given RP. We then used the mode of the MdRQA variable distributions for further analysis since it has been independently shown to be more robust to noise levels in the signal while discriminating between the states of a dynamical system with cross-validation accuracy higher than chance (Thaikkandi & Sharika, 2023). Please see Supplementary Information (Fig. S6, S7) for results comparing classifier performance using other measures of central tendency. In short, since using a sliding window size of ¡68 (¡2.5 mins) was found to yield statistically unreliable estimates of RQA variables, groups whose discussion durations were shorter than 2.5 mins (n=2) were excluded from further analysis.

The summary statistics of the different RQA variable distributions computed across the sliding windows of a given RP were used to functionally differentiate the dynamics of a set of such RPs using a nested cross-validation procedure. Nested cross validation was used to prevent the validation set from being exposed during the training stage by keeping the training set used for feature selection separate from that used as validation set for quantifying classifier performance. This was achieved by having an inner cross-validation loop to select features via best subset selection and an outer cross-validation loop to report classifier cross validation accuracy on the validation set corresponding to that outer loop.

For each iteration of the outer loop, data was divided into three parts. While two-thirds of it was used as a training set, one-third was kept as held out validation set. Best subset selection was carried out on the training set and an aggregate performance score computed for the selected combination of features using a 2-fold repeated stratified cross validation procedure. This inner loop feature selection procedure was run for 20 iterations (resulting in 40 accuracy and ROC-AUC scores) and the combination of features having the best performance score was then chosen and evaluated on the original held out validation set of the outer loop to yield the classifier performance accuracy following a 3-fold repeated stratified cross validation procedure. The outer loop was run over 200 iterations (600 iterations of 3-fold cross-validation) to construct a distribution of performance scores (cross-validation accuracy and ROC) which was plotted as box plots to assess the ability of the classifier in discriminating effective group outcomes (i.e., correct consensus or not).

## 3 Results

### 3.1 Heart rate synchrony predicts collective decisions

Heart rate data from 12 groups could not be analyzed for measures of group synchrony due to signal disruption during data acquisition for at least one group member. Overall, data from 211 participants belonging to 46 groups (7 groups with n=3, 14 groups n=4, 16 groups with n=5, 9 groups with n=6) were analyzed using custom Python implementation of Multi-dimensional Recurrence Quantification Analysis (MdRQA) (Thaikkandi & Sharika, 2023; Wallot, Roepstorff, *et al.,* 2016). The choice of MdRQA for computing a measure of synchrony was based on previous reports demonstrating the ability of non-linear methods to better account for complexity in a phenomena (say, subtle shifts in the pattern of the data owing to its sensitivity to multiple factors) when compared to traditional linear methods (Young & Benton, 2015; Perkïomäki,2011; see supplementary section A & B for details on tests done to check whether our data satisfied a sufficient condition of non-linearity, and see supplementary tables S2 & S3 for results of the tests used).

Based on the sliding window approach described in Methods (see Supplementary sections C & D for more details), only data from groups whose discussion durations lasted more than 2.5 mins were included for further analysis (n = 44). Twenty-three of 44 groups made the correct consensus decision, and 21 did not. There was no statistical difference between the group discussion durations of either set of groups (median GD duration of groups that reached the correct consensus answer = 9.3, 95% High Density Interval (HDI) = [7.7, 11]; median GD duration of groups that did not reach the correct consensus answer = 8.2, 95% HDI = [6.3, 10]), two sample Bayesian t-test gives no credible difference between the mean of these two sets of groups in terms of duration, median = -1.1, 95% HDI = [-3.6, 1.5], effect size was weak, median = -0.27, 95% HDI = [-0.89, 0.35]. The Bayes factor, *BF*_10_ = 0.41 *±* 0.01%, suggests that there is insufficient evidence to conclude for or against either hypothesis (Keysers *et al.,* 2020).

We also tested if there was a statistical difference between the group size of groups that reached a correct consensus outcome vs. those that did not and did not find that to be the case (mean size of groups that reached the correct consensus answer, median = 4.6, 95% HDI = [4.2, 5.0]; mean size of groups that did not reach the correct consensus answer, median = 4.7, 95% HDI = [4.2, 5.2]), two sample Bayesian t-test gives no credible difference between the mean of these two set of groups in terms of group size, median = 0.057, 95% HDI = [-0.59, 0.7], effect size was weak (median = -0.056, 95% HDI = [-0.58, 0.67]). The Bayes factor, *BF*_10_ = 0.3023939 *±* 0.01%, suggests that group size was not likely an important determinant of discussion outcome.

The mode of each MdRQA variable distribution across sliding windows for a given RP was z-transformed before running them through the nested cross-validated KNN classification procedure predicting whether a given group had reached correct consensus or not. We also replicated the analysis using support vector classifier (SVC) and found consistent trends (see Discussion and Supplementary Information for more details). Z-transformation was done so that the feature selection process was unbiased by differences in the absolute values of these variables. Classifier performance - cross-validation accuracy and discrimination ability (ROC AUC scores) - was compared for results obtained using time series data during the group discussion (GD) vs. different control datasets (described below) using Wilcoxon Signed rank test while ensuring that the groups being compared at each training and validation stage were the same across datasets (Dem̌sar, 2006, Cohen, 1988).

We asked if heart rate dynamics between members in each group at rest (sitting and not talking) —an index of baseline affective arousal—was sufficient to predict collective decision outcomes or whether temporal dynamics of heart rate data during group discussion was necessary. To answer this question, we ran the nested cross-validation protocol (described in Methods) separately for the following: (a) baseline (preGD) heart rate data (b) randomly shuffled heart rate data during group discussion (ran-GD) and (c) randomly shuffled heart rate data during baseline (ran-preGD). The shuffled versions were generated via random permutations to disrupt the temporal structure inherent in each times series. Comparing model performance across conditions, we found (Fig. 3) that classification accuracy was significantly higher when using MdRQA measures from group discussion compared to the control datasets: GD (median accuracy = 0.733) vs. preGD (median accuracy = 0.6) (Wilcoxon signed rank test (two sided) with Bonferroni corrected *α* = 0.0071, p = 1.923e-45, *W* =1.172e+05), vs. ran-GD (median accuracy = 0.5) (Wilcoxon signed rank test (two sided) with Bonferroni corrected *α* = 0.0071, p = 2.398e-78, *W* =1.454e+05), vs. ran-preGD (median accuracy = 0.533) (Wilcoxon signed rank test (two sided) with Bonferroni corrected *α* = 0.0071, p = 1.154e-78, *W* =1.442.0+05). The set of features that got selected most frequently during feature extraction (in the inner loop during nested cross-validation procedure, see Methods for more details) for GD were recurrence rate, percentage laminarity and average vertical line length (Fig. S3). Similarly, ROC AUC values (Fig. 4) showed significantly better classifier performance in case of GD (median ROC AUC = 0.741) vs. preGD (median ROC AUC = 0.625) (Wilcoxon signed rank test (two sided) with Bonferroni corrected *α* = 0.0071, p = 6.99e-43, *W* =1.464e+05), or ran-GD (median ROC AUC = 0.5) (Wilcoxon signed rank test (two sided) with Bonferroni corrected *α* = 0.0071, p = 3.507e-84, *W* =1.7e+05), or ran-preGD (median ROC AUC = 0.513) (Wilcoxon signed rank test (two sided) with Bonferroni corrected *α* = 0.0071, p = 4.750e-81, *W* =1.685e+05).

**Figure 4:**
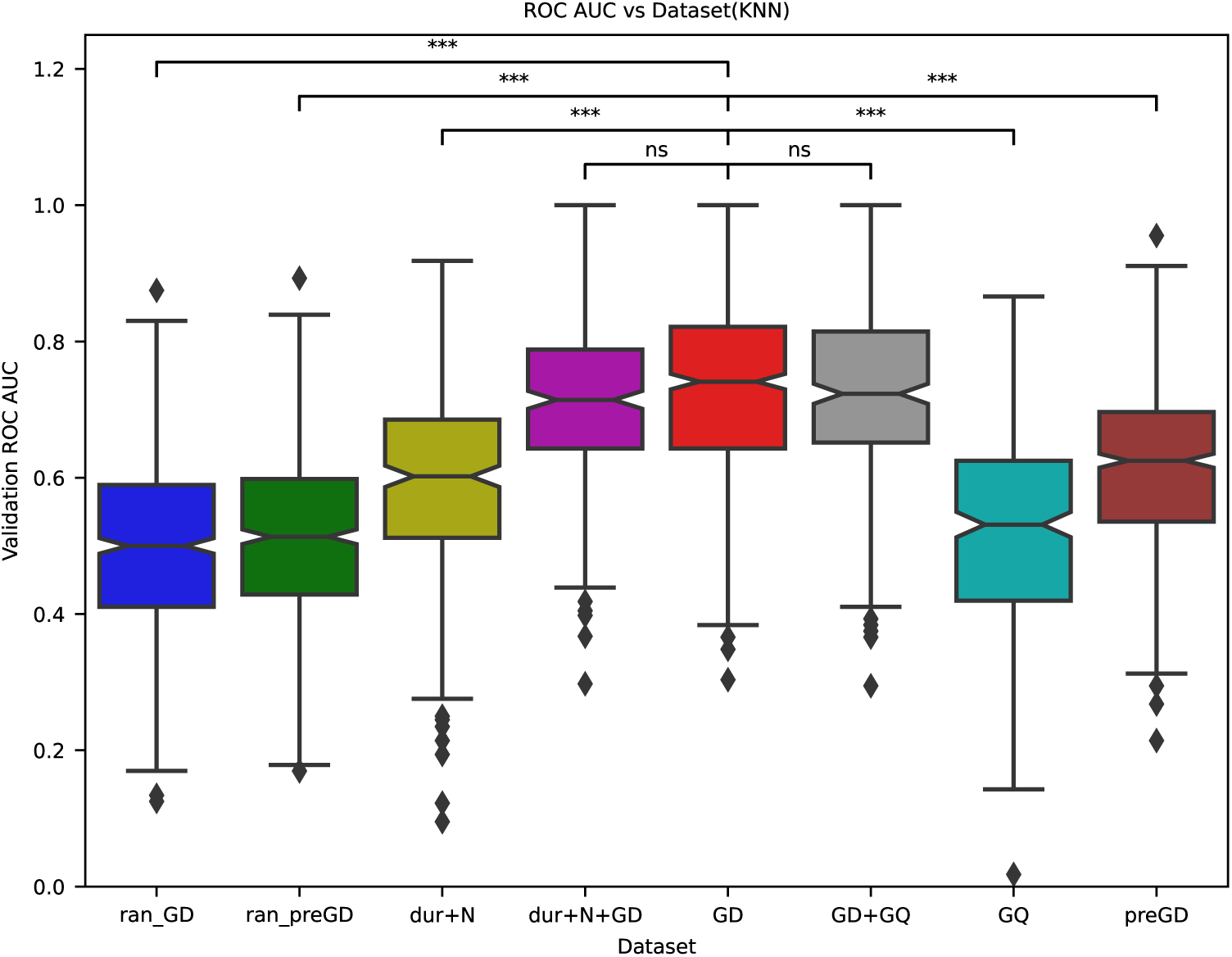
Classifier performance, represented by the area under the curve of the Receiver-Operating Characteristic Curve (ROC AUC) discriminating between groups that reached a correct consensus vs. not, was compared when it was trained using the following datasets: (a) MdRQA measures from time series data during the group discussion (GD) (b) MdRQA measures from time series data during baseline (preGD) (c) MdRQA measures from randomized time series data during group discussion (ran-GD) (d) MdRQA measures from randomized time series data during baseline (ran-preGD) (e) average group response to a triad of questionnaires on subjective group experience post-discussion (GQ) (f) MdRQA measures from time series data during the group discussion as well as the average group questionnaire response post-discussion (GD+GQ) (g) Duration (dur) and group size (N) data (dur+N) (h) MdRQA measures from time series data during the group discussion as well as duration and group size data (dur+N+GD). Identity of groups going into training and validation fold, respectively, was preserved across datasets for creating a distribution of accuracy for comparison using Wilcoxon signed-rank test. MdRQA measures refer to the mode of the MdRQA variable distribution across sliding windows of a group RP

To test how the MdRQA based heart rate synchrony measures predict effective group performance when compared to the average subjective reports of group members to questions related to the experience of working with their respective groups, we ran the KNN classification procedure with nested cross-validation using group questionnaire scores (GQ) alone as well as with MdRQA measures during group discussion (GD + GQ). We found (Fig. 3) that classification accuracy was significantly higher when using MdRQA based heart rate synchrony measures during group discussion (GD (median accuracy = 0.733) vs. when using post-discussion questionnaires scores alone (GQ (median accuracy = 0.533), Wilcoxon signed rank test (two sided) with Bonferroni corrected *α* = 0.0071, p = 9.053e-11, *W* =1.685e+05, *z* = 5.222, Cohen’s d = 0.213). Similarly, ROC AUC values (Fig. 4) showed significantly better classifier performance in case of GD (median ROC AUC = 0.732) vs. GQ (median ROC AUC = 0.533) (GD vs GQ, Wilcoxon signed rank test (two sided) with Bonferroni corrected *α* = 0.0071, p = 4.438e-36, *W* =3.497e+04). However, classification accuracy achieved using MdRQA variables from group discussion (GD (median accuracy = 0.733)) did not differ significantly when compared to that including variables from group questionnaires as well (GD + GQ (median accuracy = 0.667), Wilcoxon signed rank test (two sided) with Bonferroni corrected *α* = 0.05, p = 0.727, W=7.297e+03). The set of features that got selected most frequently during feature extraction for GD+GQ were recurrence rate, percentage laminarity, average diagonal line length and questionnaire scores from surveys on psychological safety as well as learning and well being (Fig. S7). Similarly, classifier sensitivity did not significantly differ between GD+GQ (median ROC AUC = 0.696) vs. GD (median ROC AUC = 0.732) (AUC: GD + GQ, Wilcoxon signed rank test (two sided) with Bonferroni corrected *α* = 0.05, p = 0.153, W=9.722e+03). These results suggest that while heart rate synchrony measures were significantly better at predicting group outcomes than subjective reports alone, including the latter did not help the classifier predict any better than that using heart rate data alone.

Additionally, we tested whether the group size or the duration of discussion could predict the group discussion outcome as they may indirectly affect other variables of interest. Classification accuracy for GD (median accuracy = 0.733) was significantly higher when compared to a classifier using duration and group size data alone (dur+N) (median accuracy = 0.571) (GD vs dur+N, Wilcoxon signed rank test (two sided) with Bonferroni corrected *α* = 0.0071, p = 6.243e-27, *W* =2.992e+04). Also, including these two features did not improve the accuracy of the model with the GD data (dur+N+GD) (median accuracy = 0.732)(GD vs dur+N+GD, Wilcoxon signed rank test (two sided) with Bonferroni corrected *α* = 0.0071, p = 1.964e-01, *W* =8.272e+03). Similarly, classifier sensitivity differed significantly between GD(median ROC AUC = 0.741) and classifier with duration and group size data(dur+N)(median ROC AUC = 0.602) (GD vs dur+N, Wilcoxon signed rank test (two sided) with Bonferroni corrected *α* = 0.0071, p = 6.297e-29, *W* =3.764e+04). Classifier sensitivity also did not show any improvement upon adding duration and group size as features (dur+N+GD)(GD vs dur+N+GD, Wilcoxon signed rank test (two sided) with Bonferroni corrected *α* = 0.0071, p = 3.419e-01, *W* =1.357e+04). Overall, these results suggest that group size or group discussion durations did not significantly predict group decision outcome.

## 4 Discussion

In this study, we used heart rate data collected during a naturalistic, free-flowing group-discussion task, with minimal experimental intrusion or entrainment, to ask whether interpersonal heart rate synchrony between members of a group captured the dynamics of effective decision-making. Given that non-linear methods have been reported to account for complex phenomena better than traditional linear ones (Young & Benton2015; Perkïomäki, 2011), heart rate synchrony was computed using the multi-dimensional version of recurrence quantification analysis (MdRQA) which captures the changing dynamics of multiple team members simultaneously as opposed to relying on an averaged or pair-wise measure of association (Wallot, Roepstorff, *et al.,* 2016; Thaikkandi & Sharika, 2023). Using a group discussion task based on the Hidden Profile Paradigm, where success depended on groups critically evaluating unique information against the (incorrect) option favored by shared information, we found that heart rate synchrony during group discussion, compared to pre-discussion baseline or subjective reports alone, discriminated effective groups from ineffective ones with about 70% cross-validation accuracy.

Prior work examining the role of inter-brain synchrony in group dynamics had reported its ability in predicting the overall performance of groups in a series of tasks requiring executive functioning (Reinero *et al.,* 2021). However, the inter-brain synchrony predictions were inconsistent at an individual task level. We demonstrate for the first time how even peripheral physiological markers like heart rate synchrony between members of a group larger than a dyad can predict an objectively defined criterion of group decision-making performance, (that is not based on self-reports alone) at a *single task* level with high cross-validation accuracy. This is especially noteworthy given our task did not involve, by instruction or design, participants viewing a common sensory stimulus or participating in a common motor activity as an impetus for coordinating group physiology, but rather implemented a naturalistic, group discussion scenario with high ecological validity. Though the mechanisms mediating the relationship between heart rate synchrony and this ability (or lack of) are not well-understood, prior research suggests potential underlying processes. Heart rate indexes overall autonomic arousal of an individual and, therefore, interpersonal heart rate synchrony is a proxy of shared arousal between individuals. Moreover, evidence suggests (see Palumbo *et al.,* 2017 for a detailed review) that interpersonal physiological synchrony can emerge beyond physical or metabolic demands alone (for e.g., when individuals are engaged in coordinated behaviors) and can result from psychosocial processes. For example, studies have reported increased physiological synchrony in individuals engaged in tasks demanding active reciprocity (Levenson & Gottman, 1983; Marci *et al.,* 2007; Henning *et al.,* 2001; Reed *et al.,* 2013), directed social attention or empathic accuracy (Marci, 2006; Guastello *et al.,* 2006; Richardson *et al.,* 2007; Shockley *et al.,* 2003; Schippers *et al.,* 2010; Yun *et al.,* 2012; Hasson *et al.,* 2012; Spiegelhalder *et al.,* 2014; Di Mascio *et al.,* 1955; Stratford *et al.,* 2012; Chatel-Goldman *et al.,* 2014; Marci, 2006;Ritov & Richter-Levin, 2014 ; Messina *et al.,* 2013; Jospe *et al.,* 2020). We propose that successful performance in our task taps similar mechanisms. Arriving at the correct consensus answer may require active social attention on group members during discussion to resolve differences of opinion arising from unequal distribution of information. Such motivated social engagement (Zaki, 2014), acting via processes like empathic listening & perspective-taking (Kluger & Itzchakov, 2022) may engage co-regulated physiological responses between group members, making interpersonal heart rate synchrony during discussion a strong predictor of group outcome.

Previous reports have compared group behavioral or physiological measures against to individual member measures as a way of testing whether groups outperform individuals. Owing to the nature of our decision-making task, where individual members did not have access to all the relevant information for successful performance, we did not consider making an individual vs. group performance efficacy comparison in our study. Instead, we aimed to investigate the interdependence between members’ heart rates during their interaction (also called *physiological entanglement*, Palumbo *et al.,* 2017) and its role in determining successful group performance. For this, we tested if the group synchrony measures during the interaction phase of the task (active group discussion or GD) were predictive of group efficacy, over and above what could be predicted by group synchrony measures at baseline (when group members were not interacting with each other, pre-group discussion or preGD). The randomly scrambled heart rate time series during the above two phases (ran-GD & ran-preGD, respectively) were the two additional control datasets that were used to specifically test the significance of the sequential temporal dynamics of the heart rate data in predicting group performance outcomes above chance.

Hidden profile tasks have been used in the past to understand how information transfer takes place in small groups. In a meta-analysis of 65 Hidden Profile studies, Lu et al. (Lu *et al.,* 2012) showed that groups are eight times less likely to reach the optimal decision when performing a Hidden Profile Task versus a manifest profile task, where all information is shared amongst all team members. In the context of the Hidden Profile paradigm, an increase in group size (e.g., from 3 to 8 members per group) has been typically associated with longer discussion times and greater proportion of shared information being discussed (Lu *et al.,* 2012) and ***not*** necessarily greater proportion of unique information being discussed or better group outcomes (Stasser, Taylor, *et al.,* 1989, 1992; Cruz *et al.,* 1997; Mennecke, 1997), for our purposes of testing the groups’ ability to critically evaluate alternatives by inducing them to favor the inferior option by design, groups varying from 3 to 6 members were considered to be equally amenable. The fact that multi-dimensional RQA allows accounting for different group sizes (or no. of dimensions) by keeping the recurrence rate of whole RPs constant (Wallot, Roepstorff, *et al.;* 2016) made the analysis of heart rate dynamics across the variety of group contexts (emerging out of the varying group sizes) tractable. However, as the group size and group discussion durations-based analyses showed, these features did not significantly impact the group performance outcomes in our study.

Typical behavioral biases in Hidden Profile Tasks include discussing shared information earlier and more often than unique information (Stasser & Titus, 1985; Larson Jr, 1997). This pattern of behavior is found to be sensitive to social status and group hierarchy. Specifically, lower status individuals are less likely to repeat unique information than higher status individuals, suggesting that introducing unique or dissenting information may risk social costs to individuals including negative judgement, rejection, or even exclusion from the group (Stasser & Titus, 2003, Stasser, D. D. Stewart, *et al.,* 1995; Stasser, Vaughan, *et al.,* 2000; Larson Jr, 1997; 1998, 1998; Wittenbaum, 1998; Stasser & Titus, 2003; Sohrab *et al*. 2015). In this regard, success in a Hidden Profile Task draws upon attributes previously reported to characterize effective teams, namely, psychological safety. Psychological safety describes the ability of team members to take interpersonal risks in a group by expressing, say, an unpopular opinion or unique perspective without fear of dismissal or negative evaluation (A. Edmondson, 1999; A. C. Edmondson *et al.,* 2004). In the context of our task, the ability of teams to critically examine available information and arrive at a decision contrary to the option favored by shared information would be expected to correlate with high group psychological safety. Indeed, in our data, while the classifier using heart rate synchrony measures during group discussion along with post-discussion questionnaire responses (GD + GQ) was found to perform as well as the one using heart rate data alone (GD), the top five most frequently selected features in the combined model included questionnaire scores from surveys on psychological safety as well as learning and well being (Fig. S5), hinting at the potential role of these group processes in supporting successful team performance.

It is important to note here that while we hypothesized greater heart rate synchrony would correlate with better group performance in our task, the classifier’s ability to discriminate between successful and unsuccessful group performance does not depend on any (high or low) absolute values of the RQA variables per se. Instead, among the different RQA variables derived from the RP during group discussion, recurrence rate and percentage laminarity were found to strongly contribute to the classifier’s discriminative ability (Fig. S3). This finding suggests that the overall proportion of being together as a system in a somewhat similar physiological state space as before, at multiple times during the group discussion (i.e. degree of synchrony) and for a reasonable duration at any given state (i.e. signal stability) is likely to predict successful group performance in this task.

### 4.1 Limitations

Since our final dataset had a relatively small sample size (n=44 groups), replication with a larger sample size is warranted. Increasing sample size may also support more complex statistical models such as decision trees and neural networks. k-NN as a pattern recognition technique suffers from two limitations in high dimensional applications: 1) the Euclidean distance measure suffers from the curse of dimensionality and 2) k-NN assumes all features to be equally important by default and including irrelevant features can lead to poorer predictions or generalizability on test set data. While, typically, it is datasets involving hundreds or thousands of variables as features for a sample size of less than 100 or so that are considered high dimensional exemplars (Ĺopez & Maldonado, 2018) and in our study, the number of features considered are much lower (9 to 14) for a sample size of 44, we address the concern via the nested cross-validation procedure: by first selecting for relevant features and then running the classification algorithm (Ĺopez & Maldonado, 2018). Specifically, we first ran the inner loop of cross-validation to select features in the training set (via best subset selection) and then ran the outer loop for quantifying the classifier performance in the test set. We also ran the whole analysis using an additional classification technique that is relatively more robust to high dimensionality (support vector classifier) and found overall consistent trends (see Supplementary Information, GD dataset predicted group performance outcome significantly better than GQ or other control datasets, based on both cross-validation accuracy (Fig. S6) as well as ROC AUC (Fig. S7) confirming the robustness of our results.

Heart rate data was collected from Polar chest strap sensor, H10, via their proprietary application, Polar Club, which allows temporal synchronization of data collection across all members of a group but provides already pre-processed data, as beats per minute values every 2s. Wallot, Fusaroli, *et al.,* 2013 have shown that the non-linear measures lose statistical power as the moving window size is increased from 0.3 to 6 seconds during the construction of the BPM-series from RR intervals. Though we do not have access to the exact smoothing procedure used by the application, given that we could still obtain significant differences in the predictive accuracy of GD vs. preGD (and other control datasets) using this data suggests that the relevant physiological dynamics was likely preserved and not lost by the pre-processing pipeline. However, continuous ECG data (as opposed to discrete beats per minute) could allow use of more sensitive physiological measures like heart rate variability (HRV) to better characterize physiological dynamics associated with effective group outcomes.

### 4.2 Conclusion

While prior research has linked interpersonal physiological synchrony to multiple social cognitive processes like group cohesion and empathic accuracy, whether heart rate synchrony can predict the efficacy of groups to critically evaluate alternatives, a key characteristic of information processing in group decision-making, was unknown. We addressed this gap by examining heart rate data from 44 three-to-six member groups performing a collective decision-making task based on the Hidden Profile Paradigm, in which information is distributed unequally among group members and must be evaluated critically to reach the correct consensus decision. Using multi-dimensional recurrence quantification analysis (MdRQA) and machine learning, we found that interpersonal heart rate synchrony predicted the likelihood of teams overriding inferior, default options and reaching the correct consensus decision, with more than 70 % cross-validation accuracy—significantly higher than either self-report assessment of team function or baseline heart rates prior to task onset. These findings are, to our knowledge, the first to demonstrate that heart rate synchrony provides a biomarker of effective information processing in naturalistic group decision-making.

## Acknowledgments

We thank Emily Hobart for her creative inputs on task design, Dr. Sebastien Wallot & Dr. Charles Webber for their thoughtful feedback on questions related to recurrence quantification analysis. We thank R01MH095894, R01MH108627, R37MH109728, R21AG073958, R01MH118203,866 R56MH122819 and R01NS123054, the Wharton Neuroscience Initiative and the Kishore Vaigyanik Protsahan Yojana (KVPY) scholarship programme, Department of Science and Technology, Government of India for funding the research. We also acknowledge the Wharton Behavioral Lab, University of Pennsylvania and the Remote Computing Facility at the Department of Cognitive Science, IIT Kanpur for their support.

