## supplementary information for "Interpersonal heart rate synchrony predicts effective information processing in a naturalistic group decision-making task"

March 2024

#### Abstract

Groups often outperform individuals in problem-solving. Nevertheless, failure to critically evaluate ideas risks sub-optimal outcomes through so-called *groupthink*. Prior studies have shown that people who hold shared goals, perspectives or understanding of the environment show similar patterns of brain activity, which itself can be enhanced by consensus building discussions. Whether shared arousal alone can predict collective decision-making outcomes, however, remains unknown. To address this gap, we computed interpersonal heart rate synchrony, a peripheral index of shared arousal associated with joint attention, empathic accuracy and group cohesion, in 44 groups (n=204) performing a collective decision-making task. The task required critical examination of all available information to override inferior, default options and make the right choice. Using multi-dimensional recurrence quantification analysis (MdRQA) and machine learning, we found that heart rate synchrony predicted the probability of groups reaching the correct consensus decision with greater than 70% cross-validation accuracy—significantly higher than that predicted by the duration of discussions, subjective assessment of team function or baseline heart rates alone. We propose that heart rate synchrony during group discussion provides a biomarker of interpersonal engagement that facilitates adaptive learning and effective information sharing during collective decision-making.



**Key words:** groupthink, hidden profile, heart rate synchrony, MdRQA, psychological safety

### Supplementary Video

Short Animation on the Sliding Window MdRQA Method: `https://shorturl.at/cstwT`

### Methods

#### 0.1 Test For Stationarity

Our primary objective was to see whether the time series is stationary or not, before applying a non-linear analysis method. The idea is that after entering the attractor, the time series may show stationarity, but, not when it is fairly away. We used a $\chi^2$ statistic based test to evaluate the stationarity of the time series (Isliker & Kurths, 1993). In this method we would compare the first half of the time series with the whole time series. If there is no significant difference between the two, we can consider the time series to be stationary.

If we are considering a time series of length N, we estimated Q number of bins based on the whole time series. Then we used the same bins for the first half of the signal to construct the histogram distribution. Then we compared the distribution in the first half to the distribution from the entire time series using a $\chi^2$ distribution as follows.

$$\chi^2 = \sum_{q=1}^{Q} \frac{[n_q^{half} - (\frac{n_q}{N})N_{half}]^2}{(\frac{n_q}{N})N_{half}} \tag{1}$$

here $n_q$ and $n_q^{half}$ are number of points in the $q^{th}$ bin of the histogram distribution from the whole time series and the first half respectively. We will reject the null hypothesis of stationarity if the p-value from the chi square distribution is $\leq 0.05$). We have used 10 bins for doing this. Results are given in Table **??** In total, 35 out of the 239 time series ( 15%) available was non-stationary. In other words, the data was mostly stationary.

#### 0.2 Surrogate Analysis

We tested whether the time series was non-linear, before applying RQA. Because, if the data is not non-linear, it does not necessitate the use of a non-linear method, such as RQA for analysis. We used Iterative amplitude adjusted Fourier transform (IAAFT) surrogates (Schreiber & Schmitz, 1996) for this purpose.

**Null Hypothesis:** The data represent a stationary linear Gaussian process, measured through an invertible, time-independent instantaneous measurement function(s).

Let the original signal be $x_n$, then the time independent instantaneous measurement function is given by

$$s_n = h(x_n) \tag{2}$$

Importantly

$$x_n = h^{-1}(s_n) \tag{3}$$

By linear Gaussian process, we are assuming the data represents something like the following:

$$x_n = \sum_{i=1}^{M} a_i x_{n-i} + \sum_{i=0}^{N} b_i \eta_{n-i} \tag{4}$$

Rejection of hypothesis using these surrogates suggest that the data possess some non linear structure.

The surrogate time series were created by iteratively replacing the Fourier amplitude by the correct values. A scaling is done so that the distribution and power spectrum would have close match to that of the original time series. For this we have used method and code available from (Lancaster *et al.,* 2018). The algorithm is as follows.

1. In each iteration randomly shuffle the data $y_n^{(0)}$
2. In each of the random shuffle $y_n^{(i)}$, replace the Fourier amplitudes with those from the original time series $y_n^{(0)}$. resulting in a distribution $z_n^i$ which preserve the same power spectrum but the exact distribution is not the same.
3. Rescale $z_n^i$ to original distribution of $x_n$ to produce next iteration, $y_n^{(i+1)} = x_{rank(z_n^i)}$
4. Step 2 and 3 are repeated until we reach a convergence such that $rank(z_n^i) = rank(z_n^{i+1})$ for all n.

We can either use the surrogate from step 2, which preserve the power spectrum or that from step 3 that is preserving amplitude distribution. Due to the iterative nature we cannot get a surrogate that preserve both amplitude distribution and power spectrum. And it is better to use the one which preserve power spectrum if we don't have specific hypotheses, as the rejection of null hypothesis may occur as a result of autocorrelation not being preserved in the surrogates(Kugiumtzis, 1999). As most discriminative statistics also depends on the power spectrum we decided to choose the version that preserves power spectrum. Using this method we constructed 100 surrogates for each of the time series. Then we used time irreversibility (Schreiber & Schmitz, 1997) as a measure of non-linearity, which is given by the expression:

$$\alpha = \frac{1}{N} \sum_{n=0}^{N-1} (x_{n+1} - x_n)^3 \tag{5}$$

This metric is computationally cheap to calculate as it does not require time delayed embedding. Additionally for shorter time series it maybe difficult to get a scaling region in the log-log graph between radius ($\epsilon$) and correlation sum ($C(\epsilon)$), to estimate correlation dimension. Time irreversibility indicate non-linearity as all linear systems are time reversible. But, it is important to not that time irreversibility is a sufficient, but not necessary condition for non-linearity. And this would be a two sided test. We then negated the time irreversibility measure of the original time series from each of the surrogates and performed a one sample Mann-Whitney-Wilcoxon U test to see whether the resulting distribution is significantly ($p \leq 0.05$) different from zero to see whether the time series are non-linear. Results are given in Table 3. From the results of the surrogate analysis, it is found that all time series are showing non-linearity.

Table 1: Stationarity Test Results

| Group | Member | Chi2 | DoF | p_value |
|---|---|---|---|---|
| 2 | 1 | 73.70572855 | 28 | 5.56E-06 |
| 2 | 2 | 4.990373276 | 37 | 0.9999999999 |
| 2 | 3 | 58.92545484 | 30 | 0.00123983998 |
| 2 | 4 | 13.88762288 | 29 | 0.9919314905 |
| 3 | 1 | 20.31572972 | 35 | 0.9774456993 |
| 3 | 2 | 11.89452578 | 31 | 0.9992216868 |
| 3 | 3 | 37.81622745 | 29 | 0.1264774636 |
| 3 | 4 | 19.75999767 | 24 | 0.7103394896 |
| 4 | 1 | 23.30132098 | 35 | 0.9346923666 |
| 4 | 2 | 23.77565308 | 20 | 0.2523295142 |
| 4 | 3 | 37.74323514 | 30 | 0.156453981 |
| 4 | 4 | 14.60581222 | 4 | 0.005592658505 |
| 4 | 5 | 8.416813224 | 22 | 0.9958715162 |
| 5 | 1 | 18.4619418 | 28 | 0.9139133956 |
| 5 | 2 | 18.20346887 | 23 | 0.7463610465 |
| 5 | 3 | 23.41764871 | 27 | 0.6623849798 |
| 5 | 4 | 15.35022624 | 22 | 0.8467747312 |
| 6 | 1 | 13.63490241 | 10 | 0.190305489 |
| 6 | 2 | 30.79278193 | 17 | 0.02116287817 |
| 6 | 3 | 4.27681103 | 19 | 0.9998223947 |
| 6 | 4 | 8.119047617 | 19 | 0.9854323866 |
| 6 | 5 | 3.505815884 | 13 | 0.9954030032 |
| 6 | 6 | 3.907609302 | 16 | 0.9990531413 |
| 7 | 1 | 22.91433272 | 29 | 0.7804490521 |
| 7 | 2 | 10.20983603 | 36 | 0.9999928616 |
| 7 | 3 | 24.21248212 | 32 | 0.8366232268 |
| 7 | 4 | 36.52121045 | 30 | 0.191505682 |
| 8 | 1 | 54.52797178 | 26 | 0.0008706074368 |
| 8 | 2 | 17.49639092 | 20 | 0.6205454455 |
| 8 | 3 | 32.25293413 | 36 | 0.6474892802 |
| 8 | 4 | 42.78838377 | 28 | 0.03647906107 |
| 8 | 5 | 26.44889988 | 19 | 0.1181592656 |
| 9 | 1 | 8.045843925 | 29 | 0.9999581761 |
| 9 | 2 | 16.24021479 | 23 | 0.8447941318 |

Continued on next page

**Table 1 – continued from previous page**

| Group | Member | $\chi^2$ | DoF | p_value |
|---|---|---|---|---|
| 9 | 3 | 42.9562924 | 21 | 0.003184041848 |
| 9 | 4 | 33.81625562 | 22 | 0.05127766367 |
| 9 | 5 | 6.448495992 | 20 | 0.9981401494 |
| 10 | 1 | 9.218982327 | 29 | 0.9998240829 |
| 10 | 2 | 26.97722449 | 28 | 0.5194875856 |
| 10 | 3 | 48.225554 | 15 | 2.34E-05 |
| 10 | 4 | 7.570253286 | 27 | 0.9999158689 |
| 10 | 5 | 7.619591932 | 23 | 0.9988983613 |
| 10 | 6 | 21.19863585 | 14 | 0.0966495045 |
| 11 | 1 | 21.66055892 | 35 | 0.9621526594 |
| 11 | 2 | 10.9231678 | 41 | 0.9999993338 |
| 11 | 3 | 29.61181339 | 32 | 0.5879633181 |
| 11 | 4 | 29.02731169 | 30 | 0.5161622357 |
| 11 | 5 | 4.875749128 | 26 | 0.9999981804 |
| 12 | 1 | 6.895468401 | 20 | 0.9970140439 |
| 12 | 2 | 24.90240688 | 18 | 0.1276281306 |
| 12 | 3 | 5.020903909 | 16 | 0.9956484243 |
| 14 | 1 | 13.23594654 | 35 | 0.9996844679 |
| 14 | 2 | 19.54835965 | 22 | 0.6112678522 |
| 14 | 3 | 12.16581756 | 44 | 0.9999995112 |
| 15 | 1 | 32.80421066 | 48 | 0.9538613196 |
| 15 | 2 | 15.5998204 | 41 | 0.9998888607 |
| 15 | 3 | 14.80917542 | 26 | 0.9607247802 |
| 16 | 1 | 10.84976862 | 21 | 0.9656331379 |
| 16 | 2 | 18.91451048 | 35 | 0.9878403864 |
| 16 | 3 | 35.2333144 | 30 | 0.2342500634 |
| 16 | 4 | 34.05076387 | 31 | 0.322921176 |
| 16 | 5 | 37.20490392 | 29 | 0.141055345 |
| 17 | 1 | 15.74824905 | 19 | 0.6740034064 |
| 17 | 2 | 26.24014662 | 26 | 0.4499669128 |
| 17 | 3 | 35.76893743 | 16 | 0.003114915219 |
| 17 | 4 | 20.39369368 | 19 | 0.371246804 |
| 17 | 5 | 13.63496773 | 24 | 0.9544698623 |
| 18 | 1 | 11.3323184 | 8 | 0.1835710787 |
| 18 | 2 | 0.798952984 | 7 | 0.9974546239 |
| 18 | 3 | 11.89374262 | 12 | 0.4542506686 |
| 18 | 4 | 1.14141414 | 8 | 0.997187411 |

Continued on next page

**Table 1 – continued from previous page**

| Group | Member | $\chi^2$ | DoF | p_value |
|---|---|---|---|---|
| 18 | 5 | 15.75 | 2 | 0.0003801289581 |
| 19 | 1 | 20.25629792 | 22 | 0.5670113887 |
| 19 | 2 | 19.98747211 | 33 | 0.9636388761 |
| 19 | 3 | 72.6816373 | 32 | 5.32E-05 |
| 19 | 4 | 22.21581783 | 28 | 0.7712028392 |
| 19 | 5 | 28.82813687 | 33 | 0.675028175 |
| 20 | 1 | 17.30450875 | 41 | 0.9989264799 |
| 20 | 2 | 35.08156531 | 43 | 0.7994383291 |
| 20 | 3 | 40.17015012 | 46 | 0.7140404285 |
| 20 | 4 | 16.41107092 | 38 | 0.9991288875 |
| 20 | 5 | 29.62546773 | 23 | 0.1604579491 |
| 21 | 1 | 6.31130212 | 32 | 0.9999997588 |
| 21 | 2 | 22.43515194 | 26 | 0.6646690181 |
| 21 | 3 | 23.21430751 | 19 | 0.2280343471 |
| 21 | 4 | 23.85754212 | 25 | 0.5276168215 |
| 21 | 5 | 14.15038363 | 25 | 0.9590473781 |
| 22 | 1 | 16.47636216 | 23 | 0.8341310357 |
| 22 | 2 | 25.81659249 | 25 | 0.4174374814 |
| 22 | 3 | 38.55373786 | 31 | 0.165003569 |
| 22 | 4 | 46.12246699 | 27 | 0.01234218951 |
| 22 | 5 | 35.47567509 | 31 | 0.265337246 |
| 23 | 1 | 7.903194138 | 19 | 0.9876187124 |
| 23 | 2 | 9.249599345 | 19 | 0.9691000053 |
| 23 | 3 | 10.56715892 | 19 | 0.9375417355 |
| 23 | 4 | 9.193092929 | 22 | 0.9922639712 |
| 23 | 5 | 15.44820254 | 26 | 0.9485768471 |
| 24 | 1 | 1.718800667 | 22 | 0.9999999978 |
| 24 | 2 | 11.67871548 | 16 | 0.7657846911 |
| 24 | 3 | 28.92583988 | 20 | 0.0892298883 |
| 24 | 4 | 2.623034866 | 18 | 0.9999902091 |
| 24 | 5 | 8.345583157 | 18 | 0.9730355999 |
| 25 | 1 | 13.38215661 | 16 | 0.6446372905 |
| 25 | 2 | 7.902311179 | 17 | 0.9685910592 |
| 25 | 3 | 3.658291188 | 19 | 0.9999470098 |
| 25 | 4 | 10.41724815 | 20 | 0.9599599533 |
| 25 | 5 | 15.62283127 | 14 | 0.3369458236 |
| 26 | 1 | 26.50848456 | 36 | 0.8759435027 |

Continued on next page

**Table 1 – continued from previous page**

| Group | Member | $\chi^2$ | DoF | p_value |
|---|---|---|---|---|
| 26 | 2 | 29.07310297 | 43 | 0.9484086337 |
| 26 | 3 | 30.00926474 | 24 | 0.1844449209 |
| 26 | 4 | 32.12108917 | 30 | 0.3619171616 |
| 26 | 5 | 20.20604048 | 24 | 0.6850007176 |
| 30 | 1 | 10.97124268 | 15 | 0.7546302531 |
| 30 | 2 | 16.42181818 | 15 | 0.3545845389 |
| 30 | 3 | 11.25900116 | 20 | 0.9392272215 |
| 31 | 1 | 6.873836617 | 34 | 0.9999998548 |
| 31 | 2 | 15.63087676 | 32 | 0.9933053007 |
| 31 | 3 | 10.0428531 | 35 | 0.9999889672 |
| 31 | 4 | 11.20919082 | 30 | 0.9992781885 |
| 32 | 1 | 18.02545535 | 40 | 0.9989264799 |
| 32 | 2 | 17.43425807 | 37 | 0.9974056902 |
| 32 | 3 | 12.05475895 | 40 | 0.9999944696 |
| 32 | 4 | 32.75793949 | 36 | 0.6235890582 |
| 32 | 5 | 99.67443221 | 38 | 1.96E-07 |
| 32 | 6 | 77.3444392 | 35 | 4.94E-05 |
| 33 | 1 | 10.41538995 | 14 | 0.7312270152 |
| 33 | 2 | 36.39052287 | 10 | 7.21E-05 |
| 33 | 3 | 18.55113857 | 19 | 0.485950869 |
| 33 | 4 | 7.199189236 | 18 | 0.9883363359 |
| 34 | 1 | 15.21478919 | 22 | 0.8528565019 |
| 34 | 2 | 19.55932203 | 17 | 0.2973871438 |
| 34 | 3 | 21.85949882 | 19 | 0.2913045171 |
| 34 | 4 | 35.30147058 | 9 | 5.27E-05 |
| 34 | 5 | 20.0625 | 10 | 0.0286670545 |
| 34 | 6 | 21.82590382 | 16 | 0.1489023454 |
| 35 | 1 | 20.87649448 | 20 | 0.4044338855 |
| 35 | 2 | 42.37618264 | 13 | 5.68E-05 |
| 35 | 3 | 4.264277639 | 15 | 0.9967349703 |
| 35 | 4 | 10.53408027 | 16 | 0.8372909457 |
| 36 | 1 | 11.19565688 | 14 | 0.6706017395 |
| 36 | 2 | 27.63081844 | 19 | 0.09079569988 |
| 36 | 3 | 8.355734307 | 20 | 0.9892402788 |
| 36 | 4 | 6.318597907 | 11 | 0.8512833406 |
| 36 | 5 | 24.44548577 | 17 | 0.1078273776 |
| 36 | 6 | 22.83001541 | 18 | 0.1972069964 |

Continued on next page

**Table 1 – continued from previous page**

| Group | Member | $\chi^2$ | DoF | p_value |
|---|---|---|---|---|
| 37 | 1 | 42.72827957 | 22 | 0.005096016854 |
| 37 | 2 | 12.11705139 | 31 | 0.9990628031 |
| 37 | 3 | 32.655097 | 28 | 0.2486769522 |
| 37 | 4 | 13.69630154 | 24 | 0.9532148163 |
| 38 | 1 | 8.239495797 | 17 | 0.9611539767 |
| 38 | 2 | 39.64285714 | 17 | 0.001453005338 |
| 38 | 3 | 6.153846151 | 19 | 0.9975370193 |
| 39 | 1 | 10.36046874 | 28 | 0.9990268839 |
| 39 | 2 | 13.50578745 | 19 | 0.782007044 |
| 39 | 3 | 6.664110429 | 21 | 0.9995583317 |
| 39 | 4 | 23.58536768 | 23 | 0.5069071346 |
| 39 | 5 | 4.389374552 | 17 | 0.9999963702 |
| 40 | 1 | 13.83124108 | 11 | 0.274637893 |
| 40 | 2 | 10.46777745 | 13 | 0.6420787865 |
| 40 | 3 | 13.14171295 | 11 | 0.3270566765 |
| 40 | 4 | 12.11752703 | 13 | 0.5959535935 |
| 40 | 5 | 13.60057471 | 13 | 0.4609125402 |
| 41 | 1 | 6.443644486 | 17 | 0.9899922435 |
| 41 | 2 | 2.90408805 | 22 | 0.9999999999 |
| 41 | 3 | 3.064146543 | 21 | 0.9999999769 |
| 41 | 4 | 2.810649154 | 16 | 0.9999999651 |
| 41 | 5 | 10.94430879 | 16 | 0.7445352221 |
| 42 | 1 | 14.35282258 | 13 | 0.3187640346 |
| 42 | 2 | 8.424290657 | 15 | 0.8621376353 |
| 42 | 3 | 2.802784528 | 17 | 0.9999999849 |
| 42 | 4 | 5.03802887 | 18 | 0.9999999996 |
| 42 | 5 | 10.28276452 | 14 | 0.6653899112 |
| 43 | 1 | 2.756005566 | 22 | 1 |
| 43 | 2 | 7.148260247 | 20 | 0.9718191017 |
| 43 | 3 | 5.717615943 | 21 | 0.9999999995 |
| 43 | 4 | 1.972378091 | 20 | 0.9999998822 |
| 43 | 5 | 9.591002875 | 17 | 0.860491453 |
| 44 | 1 | 2.530195927 | 21 | 1 |
| 44 | 2 | 6.651042983 | 20 | 0.9999823216 |
| 44 | 3 | 3.236747551 | 22 | 0.9999999999 |
| 44 | 4 | 4.281862459 | 21 | 0.9999999996 |
| 44 | 5 | 10.45281694 | 16 | 0.7937621196 |

Continued on next page

**Table 1 – continued from previous page**

| Group | Member | $\chi^2$ | DoF | p_value |
|---|---|---|---|---|
| 45 | 1 | 5.13274002 | 17 | 0.9999999986 |
| 45 | 2 | 3.107718438 | 20 | 0.9999999999 |
| 45 | 3 | 4.925569588 | 20 | 0.9999999993 |
| 45 | 4 | 1.274413146 | 21 | 1 |
| 45 | 5 | 8.098988372 | 14 | 0.9590707519 |
| 45 | 6 | 17.40236032 | 24 | 0.8310016893 |
| 46 | 1 | 44.14287545 | 21 | 0.002240596639 |
| 46 | 2 | 18.74621848 | 32 | 0.9697699503 |
| 46 | 3 | 14.46452146 | 29 | 0.9887602197 |
| 46 | 4 | 23.72316001 | 29 | 0.7424605644 |
| 49 | 1 | 23.85045176 | 30 | 0.7787481639 |
| 49 | 2 | 49.02964485 | 24 | 0.001878070171 |
| 49 | 3 | 48.03100246 | 21 | 0.0006809962678 |
| 49 | 4 | 19.2789434 | 27 | 0.8596141325 |
| 49 | 5 | 22.72848431 | 17 | 0.1583004084 |
| 50 | 1 | 17.23217189 | 9 | 0.04520192005 |
| 50 | 2 | 11.2847619 | 10 | 0.3357670996 |
| 50 | 3 | 19.21428571 | 8 | 0.01375471248 |
| 50 | 4 | 7.500332225 | 7 | 0.3787048959 |
| 51 | 1 | 24.7949046 | 16 | 0.07352361356 |
| 51 | 2 | 8.508065928 | 22 | 0.9955353437 |
| 51 | 3 | 18.53742966 | 19 | 0.486849436 |
| 52 | 1 | 17.18449612 | 12 | 0.1427879952 |
| 52 | 2 | 3.310030959 | 14 | 0.9983851487 |
| 52 | 3 | 20.28947368 | 19 | 0.3773520346 |
| 52 | 4 | 1.609756096 | 18 | 0.9999998102 |
| 53 | 1 | 6.218181816 | 29 | 0.9999976892 |
| 53 | 2 | 31.33605593 | 31 | 0.4493734789 |
| 53 | 3 | 8.820221088 | 22 | 0.9942181236 |
| 53 | 4 | 31.61476365 | 19 | 0.0345209755 |
| 54 | 1 | 14.96727273 | 17 | 0.5978390519 |
| 54 | 2 | 5.380519478 | 20 | 0.9995122364 |
| 54 | 3 | 4.279503104 | 16 | 0.9983317689 |
| 55 | 1 | 25.79278891 | 34 | 0.8428513878 |
| 55 | 2 | 82.68797788 | 21 | 2.86E-09 |
| 55 | 3 | 15.871994 | 31 | 0.9887758438 |
| 55 | 4 | 58.10923134 | 33 | 0.00446005007 |

Continued on next page

**Table 1 – continued from previous page**

| Group | Member | $\chi^2$ | DoF | p_value |
|---|---|---|---|---|
| 55 | 5 | 10.50047846 | 31 | 0.999786706 |
| 56 | 1 | 10.56219143 | 17 | 0.8783978621 |
| 56 | 2 | 3.795402297 | 22 | 0.9999948851 |
| 56 | 3 | 9.054901958 | 17 | 0.9385236295 |
| 56 | 4 | 5.784331796 | 22 | 0.9997847424 |
| 57 | 1 | 8.680589992 | 35 | 0.9999983779 |
| 57 | 2 | 11.52858408 | 36 | 0.9999655693 |
| 57 | 3 | 20.49584502 | 31 | 0.9245748494 |
| 57 | 4 | 17.33507082 | 37 | 0.9975569159 |
| 57 | 5 | 6.998117047 | 28 | 0.999981451 |
| 58 | 1 | 22.68421645 | 12 | 0.03052965617 |
| 58 | 2 | 24.57886588 | 13 | 0.02620430418 |
| 58 | 3 | 25.10191423 | 14 | 0.0335744992 |
| 58 | 4 | 14.33807094 | 22 | 0.8890473683 |
| 59 | 1 | 18.31941176 | 16 | 0.3055274755 |
| 59 | 2 | 4.301164675 | 18 | 0.999600307 |
| 59 | 3 | 36.83326641 | 18 | 0.005508863565 |
| 59 | 4 | 26.33481441 | 21 | 0.1939692122 |

#### 0.3 Selecting Appropriate Window Size

Recurrent plots are all those points in the trajectory such that they are within a neighbourhood, defined by radius $\epsilon$. In simple terms, let $S_n$ be the set of all points in the phase space, then the recurrent plot is given by

$$RP(i,j) = \Theta(\epsilon - ||\bar{X}_i - \bar{X}_j||) \tag{6}$$

Here it is important to note that the selection of appropriate m, $\tau$ and $\epsilon$ is important here. We choose epsilon by setting the recurrence rate to 10 percent independently on each RP.

Since we have differently sized RPs we cannot compare them directly. For this we used sliding windows with step size=1 on each RP and computed representative statistics (mean, mode and median) from these. But, the window size we have chosen should be large enough for capturing the non-linearity. For this we followed a bootstrapping method suggested in Marwan et al. (Marwan *et al.,* 2013). Let's say we have N number of windows obtained using the sliding window approach. Each of these windows will have histogram distribution of line lengths, and each window represent dynamics at a time point. Let these histogram distributions at t as a function of line length be given by $P_t(l)$. Then a unified distribution can be obtained as follows:

$$P(l) = \sum_t P_t(l) \tag{7}$$

The average number of drawings from these histogram distribution will be given as

$$\overline{n} = \frac{1}{N_{windows}} \sum_t \sum_l P_t(l) \tag{8}$$

Where the function $P_t(l)$ is actually giving the counts, and $N_{windows}$ is the total number of windows. We will now sample from the histogram distribution for getting $\overline{n}$ number of samples. The probability distribution function is given by:

$$PDF(l) = \frac{P(l)}{\sum_l P(l)} \tag{9}$$

From this we will compute the cumulative distribution function:

$$CDF(l) = \sum_{i=1}^{l} PDF(i) \tag{10}$$

Now we will random sample from this distribution by randomly drawing a number from uniform distribution [0,1] and determining to which line length that number belongs to in terms of CDF. Let the sample that we are drawing be $I_i$

$$I_i = r.v(U(0,1)) \tag{11}$$

where r.v means random variable and U(0,1) is uniform distribution between 0 and 1. Then we can apply the inverse of CDF function to get the line length that is corresponding to that particular

CDF value

$$I_i = CDF(l_i) \implies l_i = CDF^{-1}(I_i) \tag{12}$$

We repeat this $\overline{n}$ times to get one sample in the bootstrapping.

$$l = \{l_1, l_2, l_3, ..., l_{\overline{n}}\} \tag{13}$$

We will compute quantitative statistics such as mean lengths, determinism and entropy from this set. and that constitutes one bootstrapping sample.

We will repeat this sampling 1000 times to construct a distribution of nonlinear measurements for each different window size. From that we computed the difference between 95 percentile quantile and 5 percent quantile as a measure of variance, and found that the decrease is pretty low so between window size 60 and 70(see Fig 1), and based on a tolerance criteria we pick window size 68. Then we computed RQA variables using a sliding window having size 68. Provided below is the graph showing the decrease for percentage determinism. And we had to eliminate groups whose time series are shorter than 68 in terms of total number of samples.

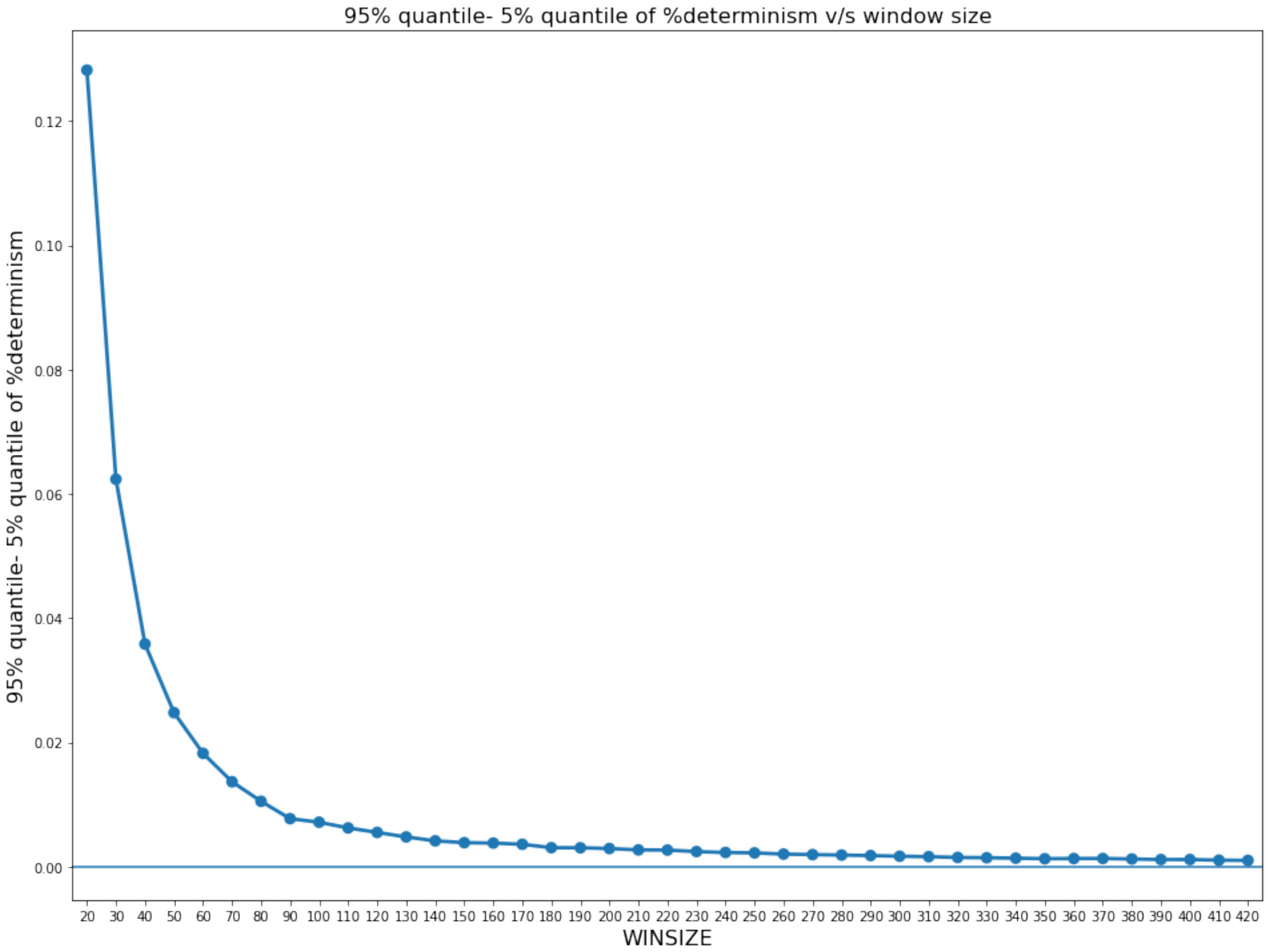


Figure 1: *The difference between 95th quantile and 5th quantile, plotted against the window size, for determinism.*

#### 0.4 RQA Parameter Estimation

##### 0.4.1 Time Delay($\tau$)

For practical purpose it is important to compute the appropriate value of the the delay($\tau$) in the first place. For this we had a multidimensional time series in which we computed a multidimensional mutual information and used it's first minima(and global minima, in case the first minima doesn't exist) in a plot between time delay and mutual information. Time delay ($\tau$) is estimated by serially sampling time delays from 1 to 20 and computing the first minima (first local minima or global minima when the local minima did not exist) of the mutual information between the time series of a group and a time delayed version of it (Wallot & Mønster, 2018). This approach ensures that the time delayed signals are not too similar and permits the multidimensional topology of the trajectories in the phase space to unfold completely. While Wallot et al (Wallot, Roepstorff, *et al.,* 2016) computed this for each time series of the group (dimension) separately and averaged the value across all members of the group, to capture the cross information between different dimensions, we computed the multidimensional mutual information by estimating the multidimensional probability distributions for the group.

##### 0.4.2 Embedding Dimension(m)

The number of embedding dimensions, $\boldsymbol{m}$, required to adequately reconstruct the phase space is estimated using the false nearest neighbor approach (Kennel *et al.,* 1992, Hegger & Kantz, 1999). False nearest neighbors (FNN) are points in the phase space that cease to be neighbors as the embedding dimension is increased and the ratio of the distances between them at the higher dimension ($\boldsymbol{m+1}$) and the current ($\boldsymbol{m}$) becomes larger than a threshold, $\boldsymbol{r}$. An increase in FNN, for a given choice of $\boldsymbol{r}$, is indicative of a phase space that needs to be reconstructed with more dimensions to unfold it completely. To arrive at an appropriate $\boldsymbol{r}$ for comparing FNN across embedding dimensions, we first plotted the FNN ratio for different values of $\boldsymbol{r}$ as embedding dimension was increased from 1 to 10. As embedding dimensions increased, we not only found FNN ratios to hit zero at smaller $\boldsymbol{r}$ values (Kantz & Schreiber, 2004, but the values of $\boldsymbol{r}$, at which FNN ratios hit zero, also varied less at the higher embedding dimensions (see Kantz & Chreiber (Kantz & Schreiber, 2004), section 3.3.1, page 37, figure 3.3) indicative of a potentially unfolded phase space. We set the tolerance criteria for the difference between the $\boldsymbol{r}$ at which FNN ratio at $\boldsymbol{m+1}$th dimension and $\boldsymbol{m}$ dimension hits zero to be 0.2 and plotted the corresponding $\boldsymbol{r}$ values as a function of embedding dimension (Supplementary Fig. 2) to select the m at which $\boldsymbol{r}$ was beginning to change the least (i.e. the knee point of the $\boldsymbol{r}$ vs. m plot- Fig. 2).

##### 0.4.3 Neighbourhood Radius($\epsilon$)

Threshold radius, $\epsilon$, which decides how close two points in the phase space should be to be considered recurrent, is chosen such that the recurrence rate (percentage of recurrent points/ black dots in the recurrence plot) is constant across different multi-component systems or RPs under study. Recurrence rate was kept constant at 10% in our case allowing RQA variables derived from different samples of a dynamical system to be directly comparable by controlling for the degree of sparseness.

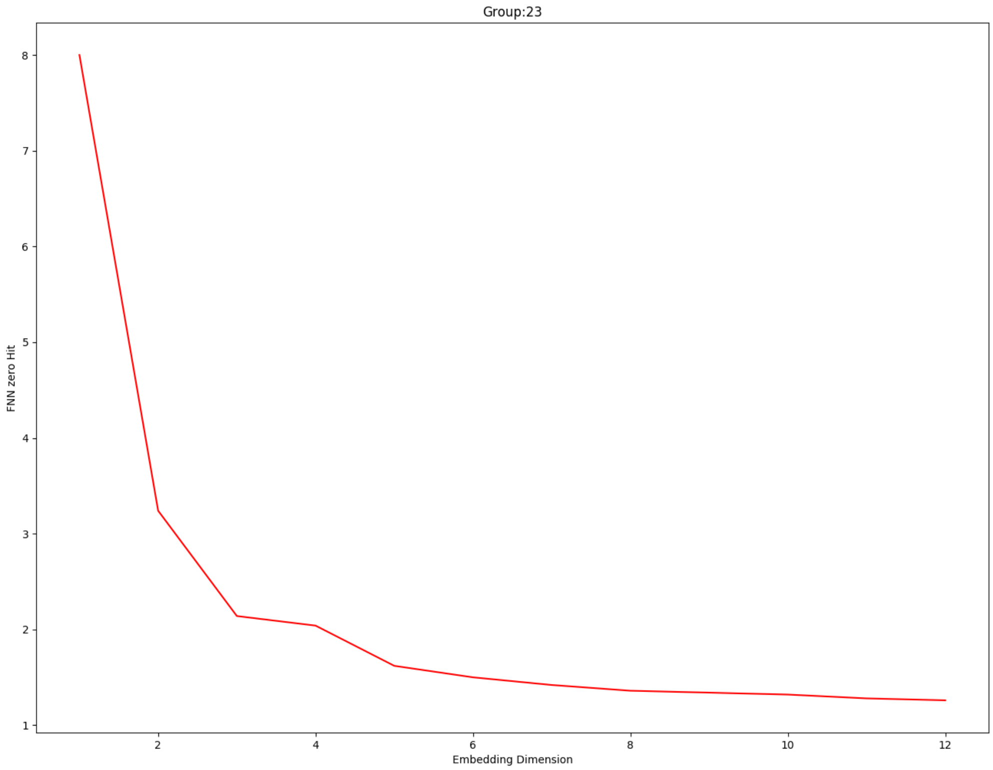


Figure 2: *FNN plotted against $r$, we can see that, as we increases the embedding dimension, the FNN hits zero for smaller values of $r$*

#### 0.5 Nested Cross Validation

**Input :** Dataset, Model information, total set of features & iterations

**Output:** Set of best features, list of performance scores on validation set

```
for each fold in the outer loop do
    Split data into training and validation sets;
    Initialize best performance score and best feature subset;
    for each feature subset(combination) do
        for each fold in inner loop do
            Split the training data further into training(train) and validation(train) sets;
            Train model on the training(train) set with feature subset;
            Calculate the performance score on the validation set for the current inner fold;
            Store the performance score for the current inner fold;
        end
        Calculate aggregate performance score for the feature subset(combination);
        if aggregate performance score is better than best performance score then
            Update best performance score;
            Update best feature subset;
        end
    end
    Store the best feature for the current outer fold;
    Train the model on training data with best feature subset for the current outer fold;
    Calculate the performance score on the validation set for the current outer fold;
    Store the performance score for the current outer fold;
end
```

**Algorithm 1:** Nested Cross-validation with Best Subset Selection

The nested cross validation is used mainly to overcome the issue of data leakage. In cross validation, we can't use selected features, as we are dividing the data multiple times into training and validation sets. To avoid data leakage, the feature selection should happen exclusively from the training set and this should be true for each of the training set division. For this, in nested cross validation, an inner loop is used within a loop that loops over all combination of features, to select features from the training set, which would be used to train model only on that training fold, which will be used to estimate the cross validation performance on the validation set for the corresponding iteration.

For each iteration of the nested CV, a feature set got selected via best subset selection. The frequency distribution of all features across all iterations quantifies how often each feature got selected. These are given in Figures 3 - 8

Table 2: Correlation matrix

| | Recurrence rate | $DET(\%)$ | $\mu_{diag}$ | $max_{diag}$ | $LAM(\%)$ | $\mu_{vert}$ | $\Sigma_{vert}$ | $\Sigma_{diag}$ | $max_{vert}$ | N | $t_{max}$ |
|---|---|---|---|---|---|---|---|---|---|---|---|
| Recurrence rate | - | - | - | - | - | - | - | - | - | - | - |
| $DET(\%)$ | -0.026 | - | ** | | * | * | * | | ** | | |
| $\mu_{diag}$ | -0.072 | 0.501 | - | | * | | | ** | | | |
| $max_{diag}$ | 0.368 | 0.345 | 0.192 | - | | *,* | | | | | * |
| $LAM(\%)$ | 0.025 | 0.635 | 0.611 | 0.251 | - | | * | | * | | |
| $\mu_{vert}$ | 0.297 | 0.279 | 0.185 | 0.411 | 0.168 | - | | | | | |
| $\Sigma_{vert}$ | 0.097 | 0.523 | 0.158 | 0.449 | 0.1 | 0.194 | - | | *** | | * |
| $\Sigma_{diag}$ | -0.035 | 0.484 | 0.503 | 0.147 | 0.432 | 0.166 | 0.269 | - | | | |
| $max_{vert}$ | 0.185 | 0.292 | -0.057 | 0.376 | 0.161 | 0.222 | 0.632 | -0.036 | - | | * |
| N | 0.253 | 0.537 | 0.278 | 0.251 | 0.402 | 0.198 | 0.293 | 0.265 | 0.148 | - | |
| $t_{max}$ | 0.278 | 0.22 | -0.171 | 0.466 | -0.072 | 0.377 | 0.425 | -0.046 | 0.475 | 0.068 | - |

#### 0.6 Comparison of Datasets

After nested cross validation, what we would have is a list of scores, and we expect using a non parametric test, such as the Wilcoxon test and Sign/Binomial test to be the best method to compare between them.

##### 0.6.1 Wilcoxon Sign Rank Test

Let, the datasets we have be A and B. Suppose, we have N iterations of a repeated cross validation and $d_i$ be the difference between the dataset A compared to B on $i^th$ iteration, given that the same statistical model with same parameters are used. Differences are ranked based on their values and average ranks are given in case of ties. Let, $R^A$ be the sum of ranks, for cases in which performance of the model on dataset A is better than that of B.

$$R^A = \sum_{d_i>0} rank(d_i) + \frac{1}{2} \sum_{d_i=0} rank(d_i) \tag{14}$$

The $R^B$ be the sum of ranks, for cases in which performance of the model on dataset B is better than that of A.

$$R^B = \sum_{d_i<0} rank(d_i) + \frac{1}{2} \sum_{d_i=0} rank(d_i) \tag{15}$$

Then, let's define $T$ as the minimum of this two sums.

$$T = \min{(R^A, R^B)} \tag{16}$$

For a large enough $N$, the z-statistic can be defined as:

$$z = \frac{T - \frac{1}{4}N(N+1)}{\sqrt{\frac{1}{24}N(N+1)(2N+1)}} \tag{17}$$

##### 0.6.2 Sign Test

Here, we are considering in each iteration, whether one dataset is performing better than the other one or not. The test statistic is simply the fraction of times $d_i$ was larger than zero.

$$P = \frac{\sum_{d_i>0} 1}{\sum_{i=1}^{N} 1} \tag{18}$$

With a large enough $N$ normal approximation can be applied to the binomial distribution, where:

$$\mu = N/2 \tag{19}$$

$$\sigma = \frac{\sqrt{N}}{2} \tag{20}$$

Depending on the hypothesis, the test can either be one sided or two sided.

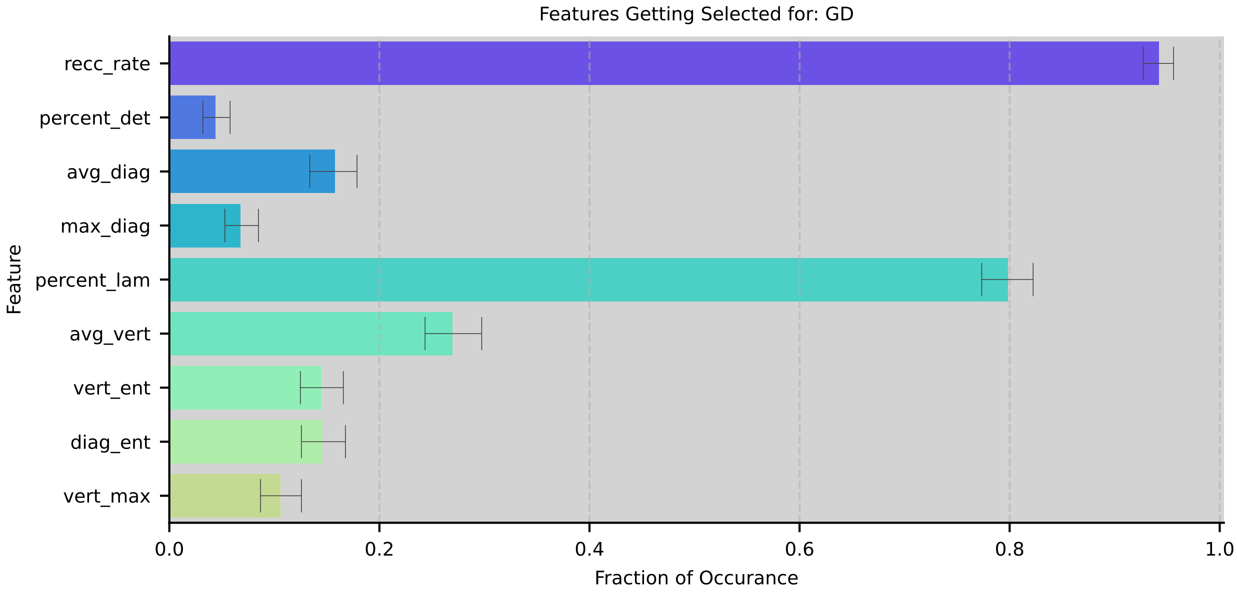


Figure 3: *Feature frequencies for GD*

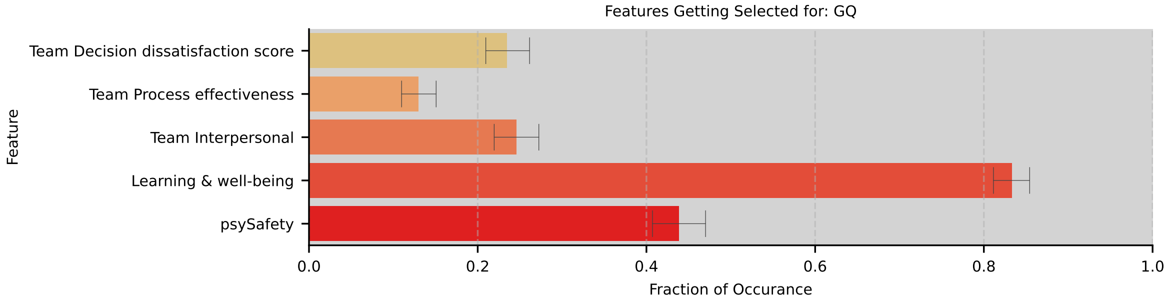


Figure 4: *Feature frequencies for GQ*

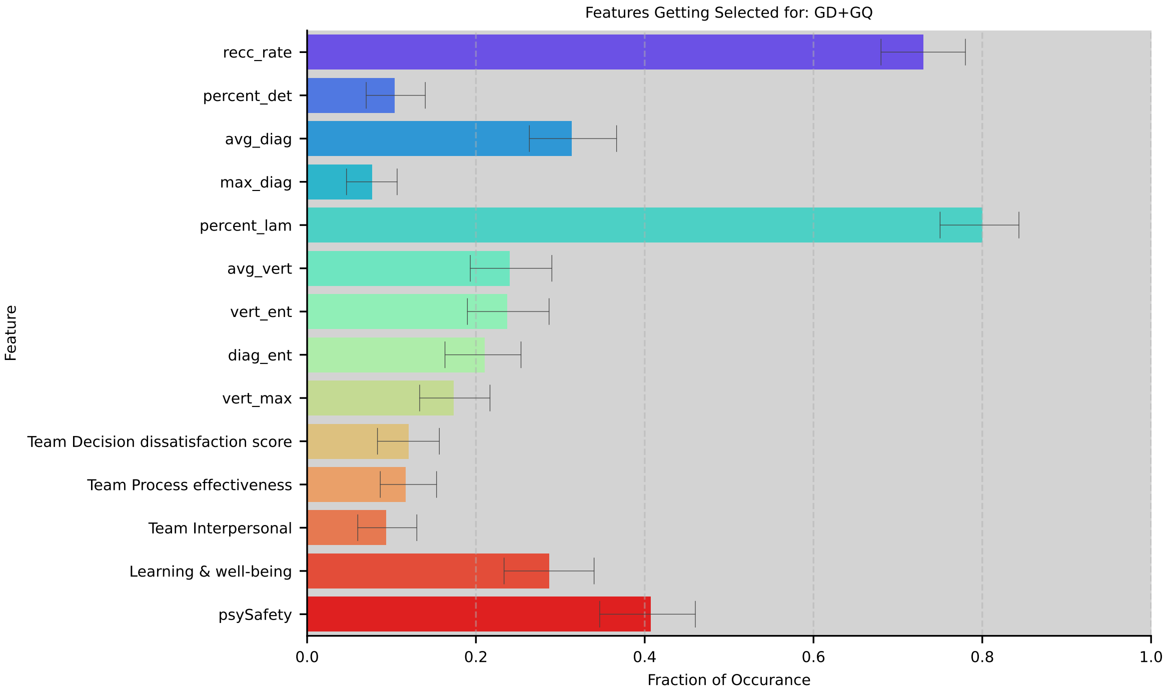


Figure 5: *Feature frequencies for GD + GQ*

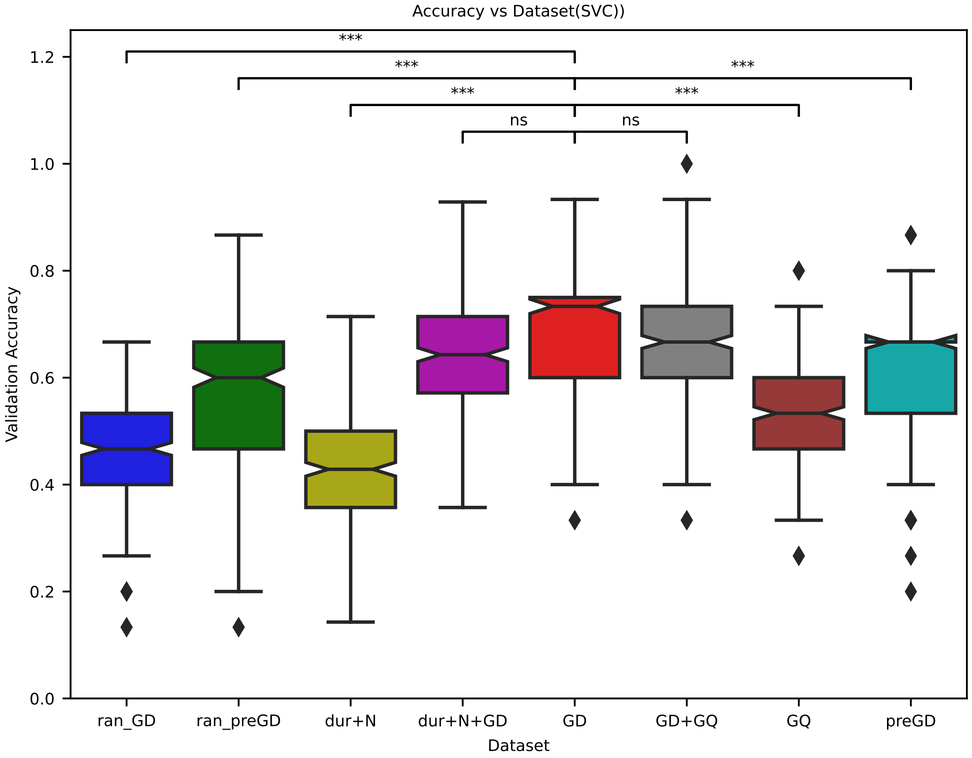


Figure 6: *Classifier performance(replication of the results using support vector classifier with linear kernel), represented by the cross-validation accuracy for discriminating between groups that reached a correct consensus vs. not, was compared when it was trained using the following datasets: (a) MdRQA measures from time series data during the group discussion (GD) (b) MdRQA measures from time series data during baseline (preGD) (c) MdRQA measures from randomized time series data during group discussion (ran-GD) (d) MdRQA measures from randomized time series data during baseline (ran-preGD) (e) average group response to a triad of questionnaires on subjective group experience post-discussion (GQ) (f) MdRQA measures from time series data during the group discussion as well as average group questionnaire response post-discussion (GD+GQ), duration and group size information (dur+N), duration and group size information with MdRQA variables. Same participants were sampled from each dataset in the training and validation set for creating a distribution of accuracy. Wilcoxon signed-rank test was then used for comparing the performance measure. MdRQA measures refer to the mode of the MdRQA variable distribution across sliding windows of a group RP. All significance levels were corrected for multiple comparison using Bon-Ferroni correction, resulting in* $\alpha = 0.0071$

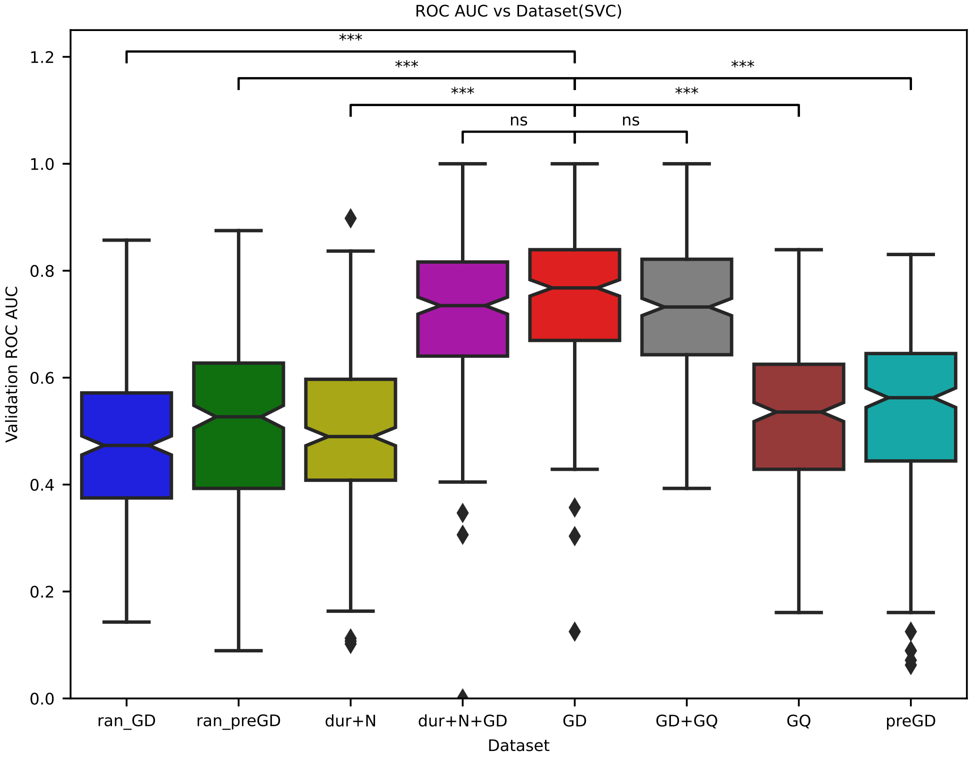


Figure 7: *Classifier performance(replication of the results using support vector classifier with linear kernel), represented by the cross-validation ROC-AUC for discriminating between groups that reached a correct consensus vs. not, was compared when it was trained using the following datasets: (a) MdRQA measures from time series data during the group discussion (GD) (b) MdRQA measures from time series data during baseline (preGD) (c) MdRQA measures from randomized time series data during group discussion (ran-GD) (d) MdRQA measures from randomized time series data during baseline (ran-preGD) (e) average group response to a triad of questionnaires on subjective group experience post-discussion (GQ) (f) MdRQA measures from time series data during the group discussion as well as average group questionnaire response post-discussion (GD+GQ), duration and group size information (dur+N), duration and group size information with MdRQA variables. Same participants were sampled from each dataset in the training and validation set for creating a distribution of accuracy. Wilcoxon signed-rank test was then used for comparing the performance measure. MdRQA measures refer to the mode of the MdRQA variable distribution across sliding windows of a group RP. All significance levels were corrected for multiple comparison using Bonferroni correction, resulting in* $\alpha$ *= 0.0071*

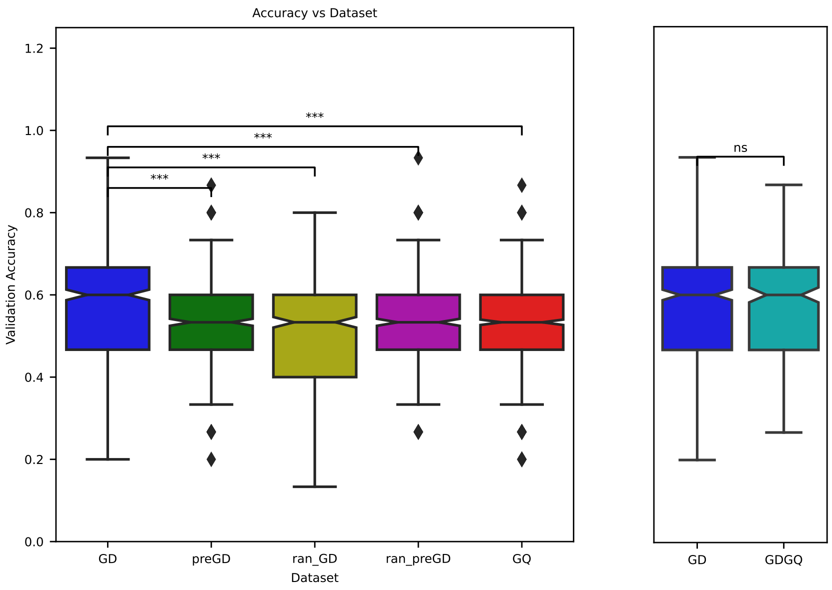


Figure 8: *Classifier performance, represented by the cross-validation accuracy for discriminating between groups that reached a correct consensus vs. not, was compared when it was trained using the following datasets: (a) MdRQA measures from time series data during the group discussion (GD) (b) MdRQA measures from time series data during baseline (preGD) (c) MdRQA measures from randomized time series data during group discussion (ran-GD) (d) MdRQA measures from randomized time series data during baseline (ran-preGD) (e) average group response to a triad of questionnaires on subjective group experience post-discussion (GQ) (f) MdRQA measures from time series data during the group discussion as well as average group questionnaire response post-discussion (GDGQ). Same participants were sampled from each dataset in the training and validation set for creating a distribution of accuracy. Wilcoxon signed-rank test was then used for comparing the performance measure. MdRQA measures refer to the mean of the MdRQA variable distribution across sliding windows of a group RP*

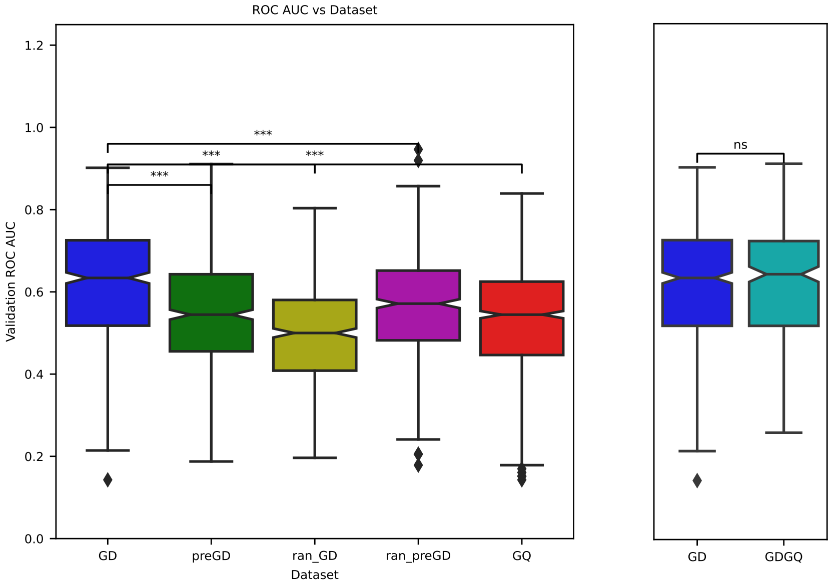


Figure 9: *Classifier performance, represented by the area under the curve of the Receiver-Operating Characteristic Curve (ROC AUC) discriminating between groups that reached a correct consensus vs. not, was compared when it was trained using the following datasets: (a) MdRQA measures from time series data during the group discussion (GD) (b) MdRQA measures from time series data during baseline (preGD) (c) MdRQA measures from randomized time series data during group discussion (ran-GD) (d) MdRQA measures from randomized time series data during baseline (ran-preGD) (e) average group response to a triad of questionnaires on subjective group experience post-discussion (GQ) (f) MdRQA measures from time series data during the group discussion as well as average group questionnaire response post-discussion (GDGQ) . Same participants were sampled from each dataset in the training and validation set for creating a distribution of accuracy. Wilcoxon signed-rank test was then used for comparing the performance measure. MdRQA measures refer to the mean of the MdRQA variable distribution across sliding windows of a group RP*

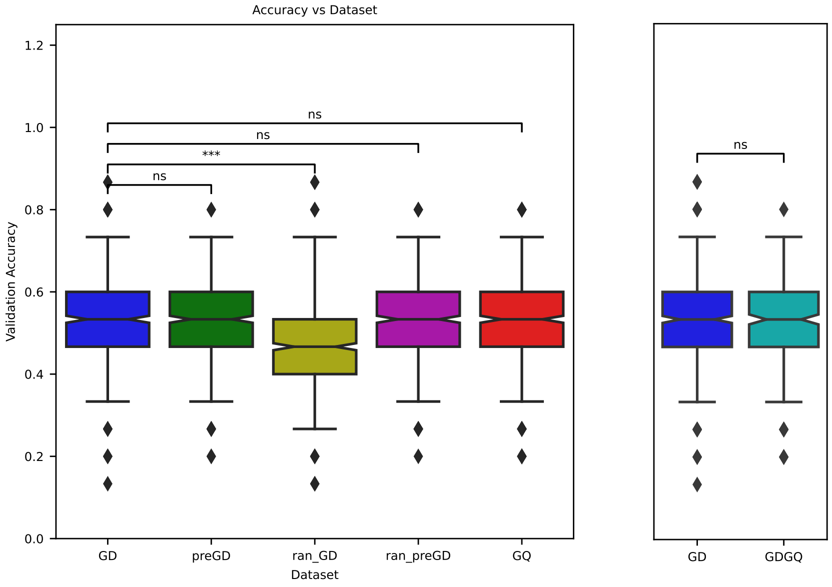


Figure 10: *Classifier performance, represented by the cross-validation accuracy for discriminating between groups that reached a correct consensus vs. not, was compared when it was trained using the following datasets: (a) MdRQA measures from time series data during the group discussion (GD) (b) MdRQA measures from time series data during baseline (preGD) (c) MdRQA measures from randomized time series data during group discussion (ran-GD) (d) MdRQA measures from randomized time series data during baseline (ran-preGD) (e) average group response to a triad of questionnaires on subjective group experience post-discussion (GQ) (f) MdRQA measures from time series data during the group discussion as well as average group questionnaire response post-discussion (GDGQ) . Same participants were sampled from each dataset in the training and validation set for creating a distribution of accuracy. Wilcoxon signed-rank test was then used for comparing the performance measure. MdRQA measures refer to the median of the MdRQA variable distribution across sliding windows of a group RP*

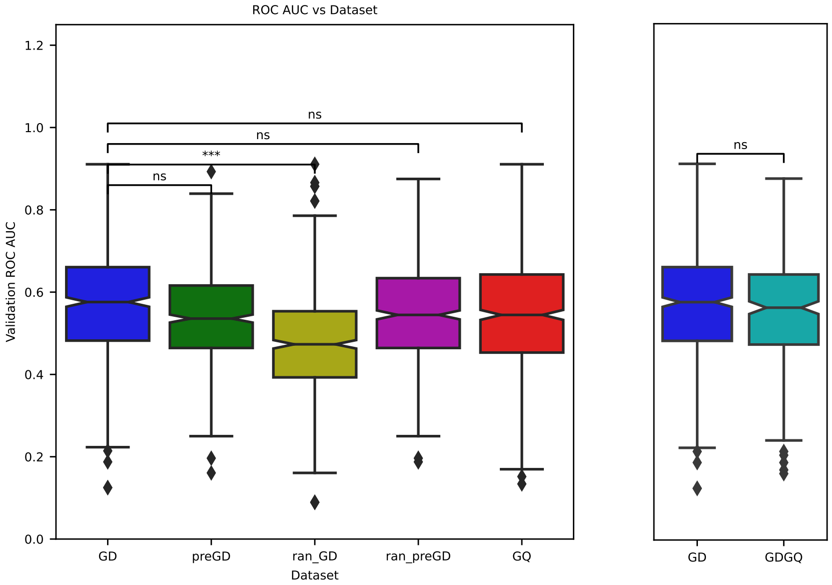


Figure 11: *Classifier performance, represented by the area under the curve of the Receiver-Operating Characteristic Curve (ROC AUC) discriminating between groups that reached a correct consensus vs. not, was compared when it was trained using the following datasets: (a) MdRQA measures from time series data during the group discussion (GD) (b) MdRQA measures from time series data during baseline (preGD) (c) MdRQA measures from randomized time series data during group discussion (ran-GD) (d) MdRQA measures from randomized time series data during baseline (ran-preGD) (e) average group response to a triad of questionnaires on subjective group experience post-discussion (GQ) (f) MdRQA measures from time series data during the group discussion as well as average group questionnaire response post-discussion (GDGQ) . Same participants were sampled from each dataset in the training and validation set for creating a distribution of accuracy. Wilcoxon signed-rank test was then used for comparing the performance measure. MdRQA measures refer to the median of the MdRQA variable distribution across sliding windows of a group RP*

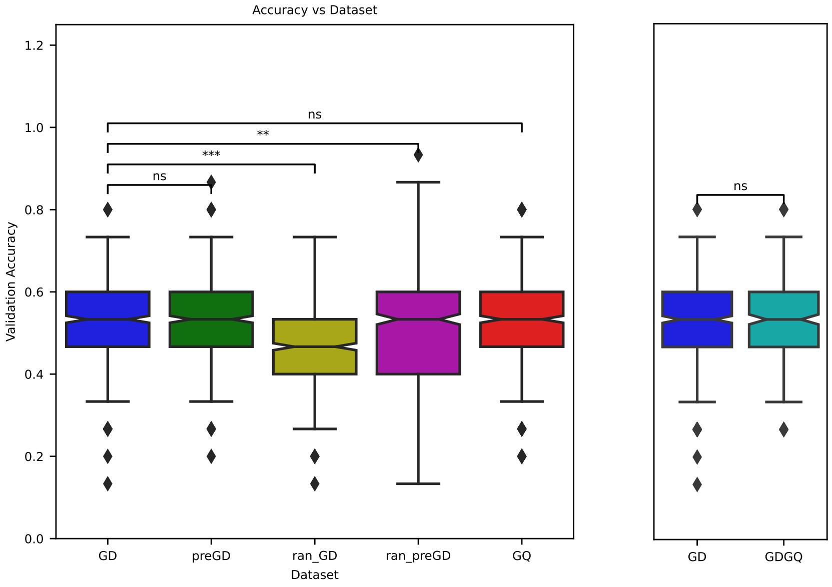


Figure 12: *Classifier performance, represented by the cross-validation accuracy for discriminating between groups that reached a correct consensus vs. not, was compared when it was trained using the following datasets: (a) MdRQA measures from time series data during the group discussion (GD) (b) MdRQA measures from time series data during baseline (preGD) (c) MdRQA measures from randomized time series data during group discussion (ran-GD) (d) MdRQA measures from randomized time series data during baseline (ran-preGD) (e) average group response to a triad of questionnaires on subjective group experience post-discussion (GQ) (f) MdRQA measures from time series data during the group discussion as well as average group questionnaire response post-discussion (GDGQ) . Same participants were sampled from each dataset in the training and validation set for creating a distribution of accuracy. Wilcoxon signed-rank test was then used for comparing the performance measure. MdRQA measures refer to variables extracted from whole RP*

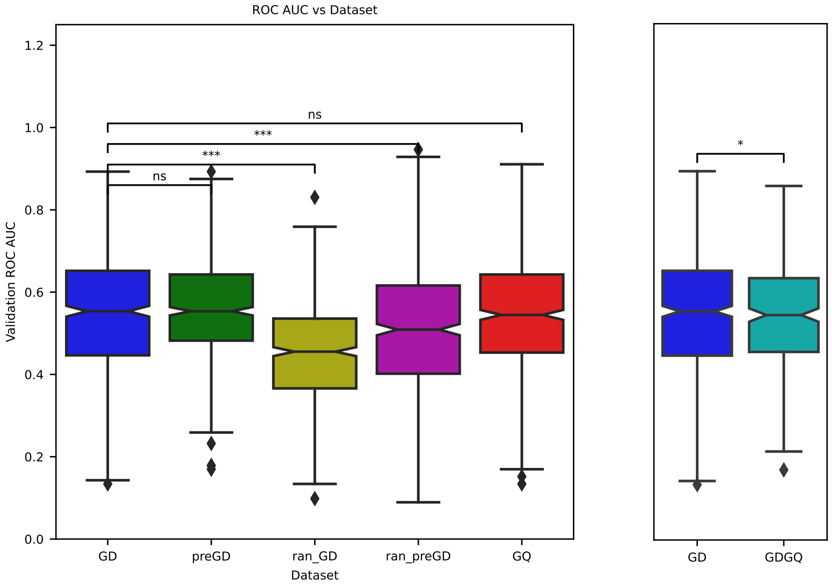


Figure 13: *Classifier performance, represented by the area under the curve of the Receiver-Operating Characteristic Curve (ROC AUC) discriminating between groups that reached a correct consensus vs. not, was compared when it was trained using the following datasets: (a) MdRQA measures from time series data during the group discussion (GD) (b) MdRQA measures from time series data during baseline (preGD) (c) MdRQA measures from randomized time series data during group discussion (ran-GD) (d) MdRQA measures from randomized time series data during baseline (ran-preGD) (e) average group response to a triad of questionnaires on subjective group experience post-discussion (GQ) (f) MdRQA measures from time series data during the group discussion as well as average group questionnaire response post-discussion (GDGQ) . Same participants were sampled from each dataset in the training and validation set for creating a distribution of accuracy. Wilcoxon signed-rank test was then used for comparing the performance measure. MdRQA measures refer to the variables extracted from whole RP*

#### 0.7 Bayesian t-test for Group Size and Discussion Duration

To test if there is any statistical difference between the distributions of group sizes and group discussion durations between groups that reached correct consensus answer vs. those that did not, we did a Bayesian t-test. Since even failing to reject the null hypothesis under test here (i.e. no difference between discussion durations of the two groups) would not allow its acceptance as per frequentist statistics, we used a Bayesian t-test. We used BayesianFirstAid (version 0.1) package in R(Kruschke, 2013). We ran an unpaired t-test, initially for the groups excluding the two groups. Additionally we computed the Bayesian evidence for the alternate hypothesis($BF_{10}$) using ttestBF function in BayesFactor package in R (Rouder *et al.,* 2009, Morey, Rouder, *et al.,* 2011, Morey & Rouder, 2011).

##### 0.7.1 For Duration Data

After eliminating groups 44 and 50, we were left with 44 groups (correct consensus = 23, incorrect consensus = 21). Firstly we ran the test, including groups 44 and 50 (correct consensus = 23, incorrect consensus = 23), and the RJAGS code under the hood is given below.

```
require(rjags)

# Setting up the data
x <- duration.incorrect
y <- duration.correct

# The model string written in the JAGS language
model_string <- "model␣{
␣␣for(i␣in␣1:length(x))␣{
␣␣␣␣x[i]␣~␣dt(␣mu_x␣,␣tau_x␣,␣nu␣)
␣␣}
␣␣x_pred␣~␣dt(␣mu_x␣,␣tau_x␣,␣nu␣)
␣␣for(i␣in␣1:length(y))␣{
␣␣␣␣y[i]␣~␣dt(␣mu_y␣,␣tau_y␣,␣nu␣)
␣␣}
␣␣y_pred␣~␣dt(␣mu_y␣,␣tau_y␣,␣nu␣)
␣␣eff_size␣<-␣(mu_x␣-␣mu_y)␣/␣sqrt((pow(sigma_x,␣2)␣+␣pow(sigma_y,␣2))␣/␣2)
␣␣mu_diff␣<-␣mu_x␣-␣mu_y
␣␣sigma_diff␣<-sigma_x␣-␣sigma_y

␣␣#␣The␣priors
␣␣mu_x␣~␣dnorm(␣mean_mu␣,␣precision_mu␣)
␣␣tau_x␣<-␣1/pow(␣sigma_x␣,␣2␣)
␣␣sigma_x␣~␣dunif(␣sigma_low␣,␣sigma_high␣)

␣␣mu_y␣~␣dnorm(␣mean_mu␣,␣precision_mu␣)
```

```
␣␣tau_y␣<-␣1/pow(␣sigma_y␣,␣2␣)
␣␣sigma_y␣~␣dunif(␣sigma_low␣,␣sigma_high␣)

␣␣#␣A␣trick␣to␣get␣an␣exponentially␣distributed␣prior␣on␣nu␣that␣starts␣at␣1.
␣␣nu␣<-␣nuMinusOne+1
␣␣nuMinusOne␣~␣dexp(1/29)
}"

# Setting parameters for the priors that in practice will result
# in flat priors on the mu's and sigma's.
mean_mu = mean( c(x, y), trim=0.2)
precision_mu = 1 / (mad( c(x, y) )^2 * 1000000)
sigma_low = mad( c(x, y) ) / 1000
sigma_high = mad( c(x, y) ) * 1000

# Initializing parameters to sensible starting values helps the convergence
# of the MCMC sampling. Here using robust estimates of the mean (trimmed)
# and standard deviation (MAD).
inits_list <- list(
  mu_x = mean(x, trim=0.2), mu_y = mean(y, trim=0.2),
  sigma_x = mad(x), sigma_y = mad(y),
  nuMinusOne = 4)

data_list <- list(
  x = x, y = y,
  mean_mu = mean_mu,
  precision_mu = precision_mu,
  sigma_low = sigma_low,
  sigma_high = sigma_high)

# The parameters to monitor.
params <- c("mu_x", "mu_y", "mu_diff", "sigma_x", "sigma_y", "sigma_diff",
            "nu", "eff_size", "x_pred", "y_pred")

# Running the model
model <- jags.model(textConnection(model_string), data = data_list,
                    inits = inits_list, n.chains = 3, n.adapt=1000)
update(model, 500) # Burning some samples to the MCMC gods....
samples <- coda.samples(model, params, n.iter=10000)

# Inspecting the posterior
plot(samples)
summary(samples)
```

#### Bayesian t-test Result for Difference In Terms of Duration Between Classes Reached Correct and Incorrect Consensus(Including group 44 & 50)

---

We provide the results below:

**Data:**

- `duration.incorrect`, $n = 23$
- `duration.correct`, $n = 23$

**Model Parameters and Generated Quantities:**

- **mu_x:** the mean of `duration.incorrect`
- **sigma_x:** the scale of `duration.incorrect`, a consistent estimate of SD when $\nu$ is large.
- **mu_y:** the mean of `duration.correct`
- **sigma_y:** the scale of `duration.correct`
- **mu_diff:** the difference in means $(\mu_x - \mu_y)$
- **sigma_diff:** the difference in scale $(\sigma_x - \sigma_y)$
- **nu:** the degrees-of-freedom for the $t$ distribution fitted to `duration.incorrect` and `duration.correct`
- **eff_size:** the effect size calculated as $\frac{\mu_x - \mu_y}{\sqrt{\frac{\sigma_x^2 + \sigma_y^2}{2}}}$
- **x_pred:** predicted distribution for a new datapoint generated as `duration.incorrect`
- **y_pred:** predicted distribution for a new datapoint generated as `duration.correct`

**Measures:**

| | mean | sd | HDIlo | HDIup | %¡comp | %¿comp |
|---|---|---|---|---|---|---|
| mu_x | 7.671 | 0.997 | 5.674 | 9.617 | 0.000 | 1.000 |
| sigma_x | 4.531 | 0.769 | 3.173 | 6.053 | 0.000 | 1.000 |
| mu_y | 9.302 | 0.832 | 7.646 | 10.901 | 0.000 | 1.000 |
| sigma_y | 3.783 | 0.652 | 2.637 | 5.112 | 0.000 | 1.000 |
| mu_diff | -1.631 | 1.292 | -4.243 | 0.829 | 0.897 | 0.103 |
| sigma_diff | 0.748 | 0.998 | -1.227 | 2.712 | 0.216 | 0.784 |
| nu | 40.588 | 30.807 | 2.980 | 101.797 | 0.000 | 1.000 |
| eff_size | -0.395 | 0.311 | -1.005 | 0.214 | 0.897 | 0.103 |
| x_pred | 7.689 | 4.936 | -1.795 | 17.572 | 0.056 | 0.944 |
| y_pred | 9.348 | 4.091 | 1.242 | 17.462 | 0.013 | 0.987 |

**Quantiles:**

| | q2.5% | q25% | median | q75% | q97.5% |
|---|---|---|---|---|---|
| mu_x | 5.699 | 7.011 | 7.669 | 8.320 | 9.648 |
| sigma_x | 3.300 | 3.995 | 4.432 | 4.968 | 6.265 |
| mu_y | 7.679 | 8.757 | 9.303 | 9.851 | 10.940 |
| sigma_y | 2.735 | 3.331 | 3.704 | 4.148 | 5.286 |
| mu_diff | -4.184 | -2.490 | -1.639 | -0.769 | 0.896 |
| sigma_diff | -1.173 | 0.105 | 0.726 | 1.370 | 2.788 |
| nu | 6.247 | 18.526 | 32.386 | 53.946 | 120.588 |
| eff_size | -1.012 | -0.602 | -0.395 | -0.184 | 0.209 |
| x_pred | -1.953 | 4.583 | 7.675 | 10.781 | 17.455 |
| y_pred | 1.292 | 6.774 | 9.321 | 11.931 | 17.536 |

See Fig. 15 for the visualization of this result.

---

#### Diagnosis for Bayesian t-test for Difference In Terms of Duration Between Classes Reached Correct and Incorrect Consensus(Including group 44 & 50)

---

Model diagnosis shows good convergence of MCMC chains. Adding the details below:

**Iterative Results:**

- Iterations: 601-10600
- Thinning interval: 1
- Number of chains: 3
- Sample size per chain: 10000

**Diagnostic Measures:**

| | mean | sd | mcmc_se | n_eff | Rhat |
|---|---|---|---|---|---|
| mu_x | 7.671 | 0.997 | 0.007 | 18452 | 1.000 |
| sigma_x | 4.531 | 0.769 | 0.007 | 12290 | 1.000 |
| mu_y | 9.302 | 0.832 | 0.006 | 17688 | 1.000 |
| sigma_y | 3.783 | 0.652 | 0.006 | 11313 | 1.002 |
| mu_diff | -1.631 | 1.292 | 0.010 | 17896 | 1.000 |
| sigma_diff | 0.748 | 0.998 | 0.009 | 12397 | 1.000 |
| nu | 40.588 | 30.807 | 0.363 | 7223 | 1.003 |
| eff_size | -0.395 | 0.311 | 0.002 | 17890 | 1.000 |
| x_pred | 7.689 | 4.936 | 0.028 | 29999 | 1.000 |
| y_pred | 9.348 | 4.091 | 0.024 | 28863 | 1.000 |

**Model Parameters and Generated Quantities:**

- **mu_x:** the mean of duration.incorrect
- **sigma_x:** the scale of duration.incorrect, a consistent estimate of SD when $\nu$ is large.
- **mu_y:** the mean of duration.correct
- **sigma_y:** the scale of duration.correct
- **mu_diff:** the difference in means $(\mu_x - \mu_y)$
- **sigma_diff:** the difference in scale $(\sigma_x - \sigma_y)$
- **nu:** the degrees-of-freedom for the $t$ distribution fitted to duration.incorrect and duration.correct
- **eff_size:** the effect size calculated as $\frac{\mu_x - \mu_y}{\sqrt{\frac{\sigma_x^2 + \sigma_y^2}{2}}}$
- **x_pred:** predicted distribution for a new datapoint generated as duration.incorrect
- **y_pred:** predicted distribution for a new datapoint generated as duration.correct

See Figures 16 & 16 for visualization of this diagnosis results.

---

#### Bayesian Factor(supporting the alternate hypothesis) Result for Difference In Terms of Duration Between Classes Reached Correct and Incorrect Consensus(Including group 44 & 50)

---

**Bayes Factor Analysis:**

- **Alternative Hypothesis:** Alt., $r = 0.707$
  - Bayes Factor: $0.6143947 \pm 0.01\%$
- **Against Denominator:** Null, $\mu_1 - \mu_2 = 0$

**Bayes Factor Type:** Independent sample

---

#### Bayesian t-test Result for Difference In Terms of Duration Between Classes Reached Correct and Incorrect Consensus(Not including group 44 & 50)

---

We provide the results below:

**Data:**

- `duration.incorrect, n = 21`
- `duration.correct, n = 23`

**Model Parameters and Generated Quantities:**

- **mu_x:** the mean of duration.incorrect
- **sigma_x:** the scale of duration.incorrect, a consistent estimate of SD when $\nu$ is large.
- **mu_y:** the mean of duration.correct
- **sigma_y:** the scale of duration.correct
- **mu_diff:** the difference in means $(\mu_x - \mu_y)$
- **sigma_diff:** the difference in scale $(\sigma_x - \sigma_y)$
- **nu:** the degrees-of-freedom for the $t$ distribution fitted to duration.incorrect and duration.correct
- **eff_size:** the effect size calculated as $\frac{\mu_x - \mu_y}{\sqrt{\frac{\sigma_x^2 + \sigma_y^2}{2}}}$
- **x_pred:** predicted distribution for a new datapoint generated as duration.incorrect
- **y_pred:** predicted distribution for a new datapoint generated as duration.correct

**Measures:**

| | mean | sd | HDIlo | HDIup | %¡comp | %¿comp |
|---|---|---|---|---|---|---|
| mu_x | 8.234 | 0.986 | 6.311 | 10.180 | 0.000 | 1.000 |
| sigma_x | 4.341 | 0.780 | 2.975 | 5.881 | 0.000 | 1.000 |
| mu_y | 9.313 | 0.827 | 7.687 | 10.941 | 0.000 | 1.000 |
| sigma_y | 3.781 | 0.639 | 2.647 | 5.095 | 0.000 | 1.000 |
| mu_diff | -1.079 | 1.287 | -3.600 | 1.470 | 0.804 | 0.196 |
| sigma_diff | 0.560 | 1.003 | -1.403 | 2.578 | 0.282 | 0.718 |
| nu | 40.666 | 30.770 | 3.330 | 101.928 | 0.000 | 1.000 |
| eff_size | -0.268 | 0.315 | -0.895 | 0.347 | 0.804 | 0.196 |
| x_pred | 8.204 | 4.729 | -1.270 | 17.458 | 0.041 | 0.959 |
| y_pred | 9.339 | 4.119 | 1.419 | 17.669 | 0.014 | 0.986 |

**Quantiles:**

| | q2.5% | q25% | median | q75% | q97.5% |
|---|---|---|---|---|---|
| mu_x | 6.308 | 7.588 | 8.229 | 8.877 | 10.179 |
| sigma_x | 3.125 | 3.801 | 4.239 | 4.767 | 6.125 |
| mu_y | 7.703 | 8.771 | 9.310 | 9.854 | 10.967 |
| sigma_y | 2.729 | 3.335 | 3.711 | 4.151 | 5.239 |
| mu_diff | -3.618 | -1.923 | -1.082 | -0.230 | 1.458 |
| sigma_diff | -1.354 | -0.087 | 0.531 | 1.172 | 2.654 |
| nu | 6.409 | 18.499 | 32.478 | 53.995 | 121.296 |
| eff_size | -0.897 | -0.476 | -0.268 | -0.057 | 0.345 |
| x_pred | -1.206 | 5.219 | 8.214 | 11.193 | 17.564 |
| y_pred | 1.259 | 6.732 | 9.329 | 11.942 | 17.556 |

See Fig. 17 for the visualization of this result.

---

### Diagnosis for Bayesian t-test for Difference In Terms of Duration Between Classes Reached Correct and Incorrect Consensus(Not including group 44 & 50)

---

Model diagnosis shows good convergence of MCMC chains. Adding the details below:

**Diagnostic Measures:**

| | mean | sd | mcmc_se | n_eff | Rhat |
|---|---|---|---|---|---|
| mu_x | 8.234 | 0.986 | 0.007 | 17746 | 1 |
| sigma_x | 4.341 | 0.780 | 0.007 | 11180 | 1 |
| mu_y | 9.313 | 0.827 | 0.006 | 17853 | 1 |
| sigma_y | 3.781 | 0.639 | 0.006 | 13331 | 1 |
| mu_diff | -1.079 | 1.287 | 0.010 | 18046 | 1 |
| sigma_diff | 0.560 | 1.003 | 0.009 | 12529 | 1 |
| nu | 40.666 | 30.770 | 0.385 | 6517 | 1 |
| eff_size | -0.268 | 0.315 | 0.002 | 18369 | 1 |
| x_pred | 8.204 | 4.729 | 0.028 | 29549 | 1 |
| y_pred | 9.339 | 4.119 | 0.024 | 29671 | 1 |

**MCMC Diagnostics:**

- **mcmc_se:** the estimated standard error of the MCMC approximation of the mean.
- **n_eff:** a crude measure of effective MCMC sample size.
- **Rhat:** the potential scale reduction factor (at convergence, Rhat=1).

**Model Parameters and Generated Quantities:**

- **mu_x:** the mean of duration.incorrect
- **sigma_x:** the scale of duration.incorrect, a consistent estimate of SD when $\nu$ is large.
- **mu_y:** the mean of duration.correct
- **sigma_y:** the scale of duration.correct
- **mu_diff:** the difference in means $(\mu_x - \mu_y)$
- **sigma_diff:** the difference in scale $(\sigma_x - \sigma_y)$
- **nu:** the degrees-of-freedom for the $t$ distribution fitted to duration.incorrect and duration.correct
- **eff_size:** the effect size calculated as $\frac{\mu_x - \mu_y}{\sqrt{\frac{\sigma_x^2 + \sigma_y^2}{2}}}$
- **x_pred:** predicted distribution for a new datapoint generated as duration.incorrect
- **y_pred:** predicted distribution for a new datapoint generated as duration.correct

See Figures 18 & 18 for visualization of this diagnosis results.

##### 0.7.2 For Group Size Data

Here, we are treating the group size(number of members) as a continuous variable for running the Bayesian t-test.

---

#### Bayesian Factor(supporting the alternate hypothesis) Result for Difference In Terms of Duration Between Classes Reached Correct and Incorrect Consensus(Not including group 44 & 50)

---

**Bayes Factor Analysis:**

- **Alternative Hypothesis:** Alt., $r = 0.707$
  - Bayes Factor: $0.4120457 \pm 0.01\%$
- **Against Denominator:** Null, $\mu_1 - \mu_2 = 0$

**Bayes Factor Type:** Independent sample

---

#### Bayesian t-test Result for Difference In Terms of Group Size(N) Between Classes Reached Correct and Incorrect Consensus(Including group 44 & 50)

Results are provided below:

**Data:**

| | |
|---|---|
| N.incorrect | $n = 23$ |
| N.correct | $n = 23$ |

**Model Parameters and Generated Quantities:**

- **mu_x:** the mean of N.incorrect
- **sigma_x:** the scale of N.incorrect, a consistent estimate of SD when $\nu$ is large.
- **mu_y:** the mean of N.correct
- **sigma_y:** the scale of N.correct
- **mu_diff:** the difference in means $(\mu_x - \mu_y)$
- **sigma_diff:** the difference in scale $(\sigma_x - \sigma_y)$
- **nu:** the degrees-of-freedom for the $t$ distribution fitted to N.incorrect and N.correct
- **eff_size:** the effect size calculated as $\frac{\mu_x - \mu_y}{\sqrt{\frac{\sigma_x^2 + \sigma_y^2}{2}}}$
- **x_pred:** predicted distribution for a new datapoint generated as N.incorrect
- **y_pred:** predicted distribution for a new datapoint generated as N.correct

**Measures:**

| | mean | sd | HDIlo | HDIup | %¡comp | %¿comp |
|---|---|---|---|---|---|---|
| mu_x | 4.565 | 0.236 | 4.100 | 5.035 | 0.000 | 1.000 |
| sigma_x | 1.078 | 0.180 | 0.764 | 1.450 | 0.000 | 1.000 |
| mu_y | 4.613 | 0.214 | 4.192 | 5.033 | 0.000 | 1.000 |
| sigma_y | 0.976 | 0.168 | 0.674 | 1.314 | 0.000 | 1.000 |
| mu_diff | -0.047 | 0.319 | -0.656 | 0.597 | 0.562 | 0.438 |
| sigma_diff | 0.102 | 0.245 | -0.390 | 0.582 | 0.330 | 0.670 |
| nu | 41.101 | 30.629 | 2.766 | 102.219 | 0.000 | 1.000 |
| eff_size | -0.047 | 0.307 | -0.652 | 0.553 | 0.562 | 0.438 |
| x_pred | 4.565 | 1.173 | 2.204 | 6.856 | 0.001 | 0.999 |
| y_pred | 4.620 | 1.060 | 2.520 | 6.705 | 0.000 | 1.000 |

**Quantiles:**

| | q2.5% | q25% | median | q75% | q97.5% |
|---|---|---|---|---|---|
| mu_x | 4.094 | 4.411 | 4.565 | 4.720 | 5.031 |
| sigma_x | 0.789 | 0.950 | 1.056 | 1.180 | 1.496 |
| mu_y | 4.187 | 4.474 | 4.613 | 4.752 | 5.031 |
| sigma_y | 0.705 | 0.858 | 0.955 | 1.071 | 1.363 |
| mu_diff | -0.673 | -0.257 | -0.049 | 0.164 | 0.581 |
| sigma_diff | -0.377 | -0.055 | 0.101 | 0.254 | 0.599 |
| nu | 6.489 | 19.021 | 33.086 | 54.653 | 120.859 |
| eff_size | -0.651 | -0.252 | -0.048 | 0.161 | 0.556 |
| x_pred | 2.231 | 3.818 | 4.559 | 5.315 | 6.893 |
| y_pred | 2.527 | 3.948 | 4.619 | 5.289 | 6.720 |

See Fig. 19 for visualization of this result.

---

### Diagnosis for Bayesian t-test for Difference In Terms of Group Size Between Classes Reached Correct and Incorrect Consensus(Including group 44 & 50)

---

**MCMC Diagnostic Measures:**

| | mean | sd | mcmc_se | n_eff | Rhat |
|---|---|---|---|---|---|
| mu_x | 4.565 | 0.236 | 0.002 | 18717 | 1.000 |
| sigma_x | 1.078 | 0.180 | 0.002 | 13221 | 1.000 |
| mu_y | 4.613 | 0.214 | 0.002 | 17321 | 1.000 |
| sigma_y | 0.976 | 0.168 | 0.001 | 12555 | 1.000 |
| mu_diff | -0.047 | 0.319 | 0.002 | 18122 | 1.000 |
| sigma_diff | 0.102 | 0.245 | 0.002 | 13530 | 1.000 |
| nu | 41.101 | 30.629 | 0.375 | 6945 | 1.002 |
| eff_size | -0.047 | 0.307 | 0.002 | 18431 | 1.000 |
| x_pred | 4.565 | 1.173 | 0.007 | 29999 | 1.000 |
| y_pred | 4.620 | 1.060 | 0.006 | 28943 | 1.000 |

**MCMC Se, Effective Sample Size, and Potential Scale Reduction Factor:**

- **mcmc_se:** the estimated standard error of the MCMC approximation of the mean.
- **n_eff:** a crude measure of effective MCMC sample size.
- **Rhat:** the potential scale reduction factor (at convergence, Rhat=1).

**Model Parameters and Generated Quantities:**

- **mu_x:** the mean of N.incorrect

- **sigma_x:** the scale of N.incorrect, a consistent estimate of SD when $\nu$ is large.
- **mu_y:** the mean of N.correct
- **sigma_y:** the scale of N.correct
- **mu_diff:** the difference in means $(\mu_x - \mu_y)$
- **sigma_diff:** the difference in scale $(\sigma_x - \sigma_y)$
- **nu:** the degrees-of-freedom for the $t$ distribution fitted to N.incorrect and N.correct
- **eff_size:** the effect size calculated as $\frac{\mu_x - \mu_y}{\sqrt{\frac{\sigma_x^2 + \sigma_y^2}{2}}}$
- **x_pred:** predicted distribution for a new datapoint generated as N.incorrect
- **y_pred:** predicted distribution for a new datapoint generated as N.correct

See Figures 20 & 20 for visualization of this diagnosis results.

### Bayesian Factor(supporting the alternate hypothesis) Result for Difference In Terms of Group Size(N) Between Classes Reached Correct and Incorrect Consensus(Including group 44 & 50)

---

**Bayes Factor Analysis:**

- **Alternative Hypothesis:** Alt., $r = 0.707$
  - Bayes Factor: $0.2949192 \pm 0.01\%$
- **Against Denominator:** Null, $\mu_1 - \mu_2 = 0$

**Bayes Factor Type:** Independent sample

---

### Bayesian t-test Result for Difference In Terms of Group Size(N) Between Classes Reached Correct and Incorrect Consensus(Not including group 44 & 50)

---

Results are provided below:

| | N.incorrect | N.correct |
|---|---|---|
| $n$ | 21 | 23 |

**Model Parameters and Generated Quantities:**

- **mu_x:** the mean of N.incorrect
- **sigma_x:** the scale of N.incorrect, a consistent estimate of SD when $\nu$ is large.
- **mu_y:** the mean of N.correct
- **sigma_y:** the scale of N.correct
- **mu_diff:** the difference in means $(\mu_x - \mu_y)$
- **sigma_diff:** the difference in scale $(\sigma_x - \sigma_y)$
- **nu:** the degrees-of-freedom for the $t$ distribution fitted to N.incorrect and N.correct
- **eff_size:** the effect size calculated as $\frac{\mu_x - \mu_y}{\sqrt{\frac{\sigma_x^2 + \sigma_y^2}{2}}}$
- **x_pred:** predicted distribution for a new datapoint generated as N.incorrect
- **y_pred:** predicted distribution for a new datapoint generated as N.correct

**Measures:**

| | mean | sd | HDIlo | HDIup | %¡comp | %¿comp |
|---|---|---|---|---|---|---|
| mu_x | 4.670 | 0.247 | 4.173 | 5.152 | 0.000 | 1.000 |
| sigma_x | 1.064 | 0.191 | 0.729 | 1.442 | 0.000 | 1.000 |
| mu_y | 4.613 | 0.213 | 4.198 | 5.045 | 0.000 | 1.000 |
| sigma_y | 0.973 | 0.168 | 0.674 | 1.313 | 0.000 | 1.000 |
| mu_diff | 0.057 | 0.328 | -0.590 | 0.700 | 0.431 | 0.569 |
| sigma_diff | 0.091 | 0.253 | -0.413 | 0.588 | 0.355 | 0.645 |
| nu | 39.895 | 30.950 | 2.896 | 100.579 | 0.000 | 1.000 |
| eff_size | 0.057 | 0.318 | -0.577 | 0.669 | 0.431 | 0.569 |
| x_pred | 4.676 | 1.152 | 2.393 | 6.949 | 0.000 | 1.000 |
| y_pred | 4.598 | 1.063 | 2.469 | 6.693 | 0.000 | 1.000 |

**Quantiles:**

| | q2.5% | q25% | median | q75% | q97.5% |
|---|---|---|---|---|---|
| mu_x | 4.179 | 4.507 | 4.670 | 4.831 | 5.161 |
| sigma_x | 0.761 | 0.929 | 1.041 | 1.172 | 1.500 |
| mu_y | 4.188 | 4.475 | 4.612 | 4.751 | 5.036 |
| sigma_y | 0.704 | 0.855 | 0.952 | 1.068 | 1.361 |
| mu_diff | -0.586 | -0.161 | 0.057 | 0.273 | 0.706 |
| sigma_diff | -0.402 | -0.070 | 0.087 | 0.247 | 0.603 |
| nu | 6.263 | 18.020 | 31.521 | 52.546 | 121.841 |
| eff_size | -0.563 | -0.158 | 0.056 | 0.269 | 0.684 |
| x_pred | 2.396 | 3.950 | 4.673 | 5.407 | 6.954 |
| y_pred | 2.477 | 3.934 | 4.595 | 5.269 | 6.706 |

See Fig. 21 for visualization of this result.

---

### Diagnosis for Bayesian t-test for Difference In Terms of Group Size Between Classes Reached Correct and Incorrect Consensus(Not including group 44 & 50)

---

**Model Summary:**

- **Iterations:** 601–10600
- **Thinning interval:** 1
- **Number of chains:** 3
- **Sample size per chain:** 10000

**Diagnostic Measures:**

| | mean | sd | mcmc_se | n_eff | Rhat |
|---|---|---|---|---|---|
| mu_x | 4.670 | 0.247 | 0.002 | 18387 | 1.001 |
| sigma_x | 1.064 | 0.191 | 0.002 | 12162 | 1.000 |
| mu_y | 4.613 | 0.213 | 0.002 | 18442 | 1.000 |
| sigma_y | 0.973 | 0.168 | 0.001 | 12891 | 1.000 |
| mu_diff | 0.057 | 0.328 | 0.002 | 17899 | 1.000 |
| sigma_diff | 0.091 | 0.253 | 0.002 | 12581 | 1.000 |
| nu | 39.895 | 30.950 | 0.389 | 6373 | 1.006 |
| eff_size | 0.057 | 0.318 | 0.002 | 18356 | 1.000 |
| x_pred | 4.676 | 1.152 | 0.007 | 29580 | 1.000 |
| y_pred | 4.598 | 1.063 | 0.006 | 28579 | 1.000 |

**Model Parameters and Generated Quantities:**

- **mu_x:** the mean of N.incorrect
- **sigma_x:** the scale of N.incorrect, a consistent estimate of SD when $\nu$ is large.
- **mu_y:** the mean of N.correct
- **sigma_y:** the scale of N.correct
- **mu_diff:** the difference in means $(\mu_x - \mu_y)$
- **sigma_diff:** the difference in scale $(\sigma_x - \sigma_y)$
- **nu:** the degrees-of-freedom for the $t$ distribution fitted to N.incorrect and N.correct

- **eff_size:** the effect size calculated as $\frac{\mu_x - \mu_y}{\sqrt{\frac{\sigma_x^2 + \sigma_y^2}{2}}}$
- **x_pred:** predicted distribution for a new datapoint generated as N.incorrect
- **y_pred:** predicted distribution for a new datapoint generated as N.correct

**MCMC Summary:**

- **mcmc_se:** the estimated standard error of the MCMC approximation of the mean.
- **n_eff:** a crude measure of effective MCMC sample size.
- **Rhat:** the potential scale reduction factor (at convergence, Rhat=1).

See Figures 22 & 22 for visualization of this diagnosis results.

---

### Bayesian Factor(supporting the alternate hypothesis) Result for Difference In Terms of Group Size(N) Between Classes Reached Correct and Incorrect Consensus(Not including group 44 & 50)

---

**Bayes Factor Analysis:**

- **Alternative Hypothesis:** Alt., $r = 0.707$
  - Bayes Factor: $0.3023939 \pm 0.01\%$
- **Against Denominator:** Null, $\mu_1 - \mu_2 = 0$

**Bayes Factor Type:** Independent sample

After including the two groups excluded due to short discussion durations: GD duration of groups that reached the correct consensus answer, median = 9.3, 95% HDI = [7.7, 11]; GD duration of groups that did not reach the correct consensus answer, median = 7.7, 95% HDI = [5.7, 9.6]), two sample Bayesian t-test gives no credible difference between the mean of these two sets of groups in terms of duration, median = -1.6, 95% HDI = [-4.2, 0.83], effect size was weak, median = -0.39, 95% HDI = [-1.0, 0.21], Bayesian evidence for the alternate hypothesis was very weak, $BF_{10} = 0.61 \pm 0.01\%$.

After including the two groups excluded for having short discussion durations, mean size of groups that reached the correct consensus answer, median = 4.6, 95% HDI = [4.2, 5.0]; mean size of groups that did not reach the correct consensus answer, median = 4.6, 95% HDI = [4.1, 5.0]), two sample Bayesian t-test gives no credible difference between the mean of these two set of groups in terms of mean group size, median = -0.049, 95% HDI = [-0.66, 0.60], effect size was weak, median = -0.048, 95% HDI = [-0.65, 0.55]. Bayesian evidence for the alternate hypothesis was very weak,$BF_{10} = 0.2949192 \pm 0.01\%$.

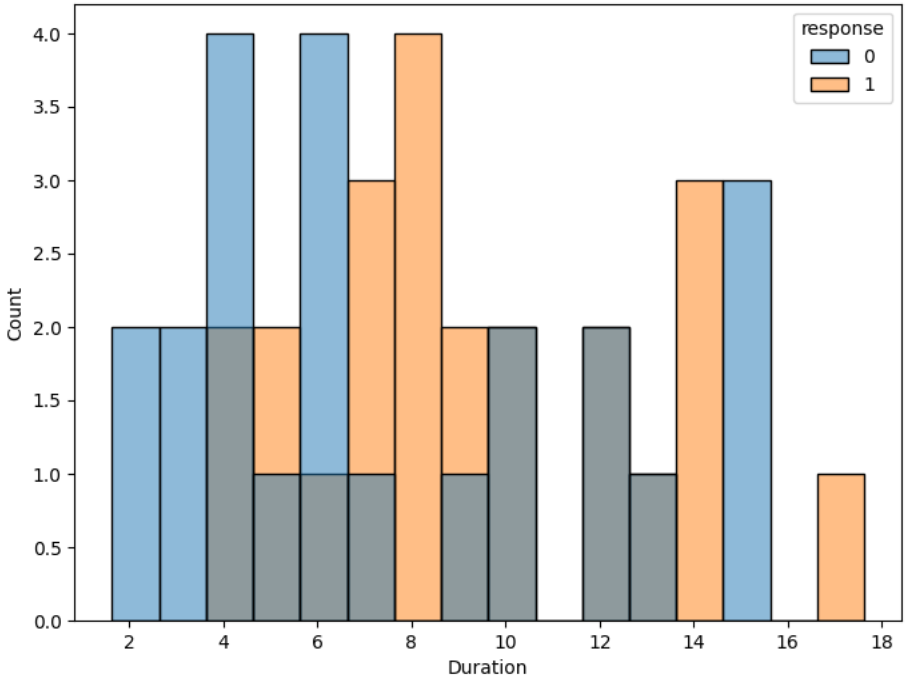


Figure 14: *Distributions of discussion durations for groups that reached correct consensus(response = 1) and did not reached correct consensus(response = 0)*

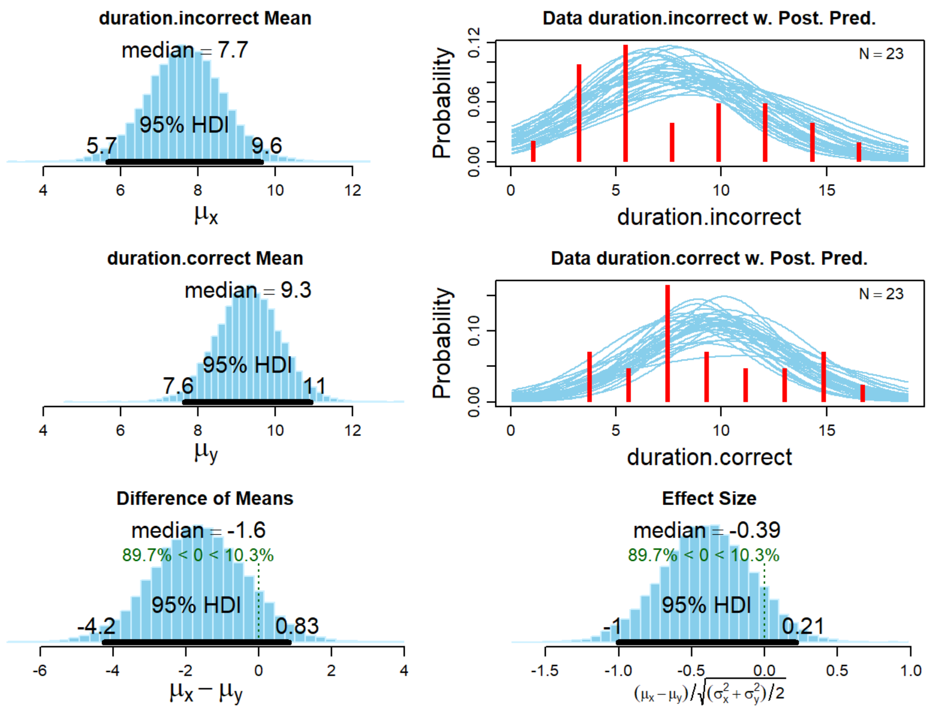


Figure 15: *Bayesian t-test results for the duration data, when including the excluded groups (groups 44 & 50 due to short duration)*

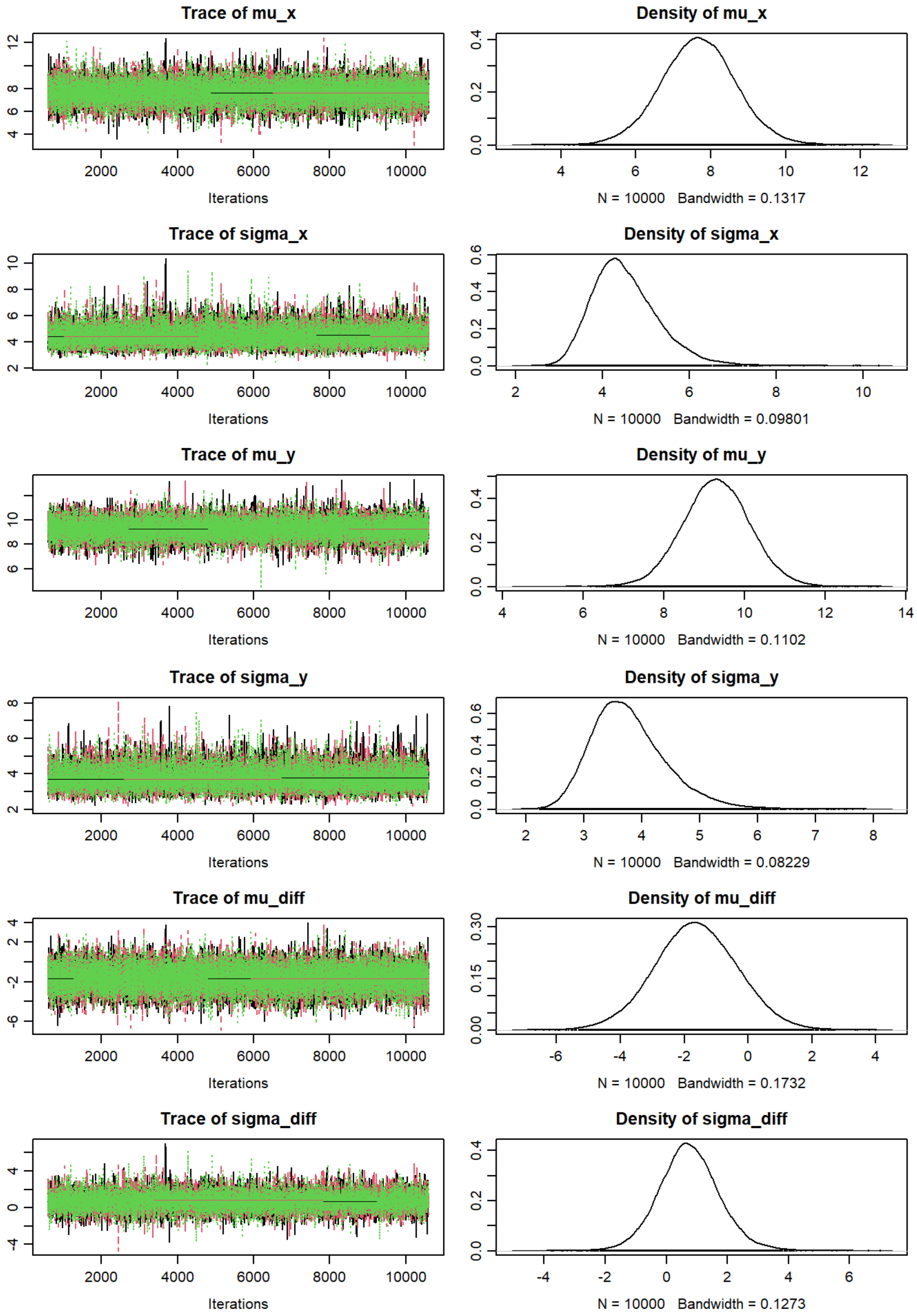


Figure 16: *Bayesian t-test diagnosis plots for the duration data, when including the excluded groups (groups 44 & 50 due to short duration)*

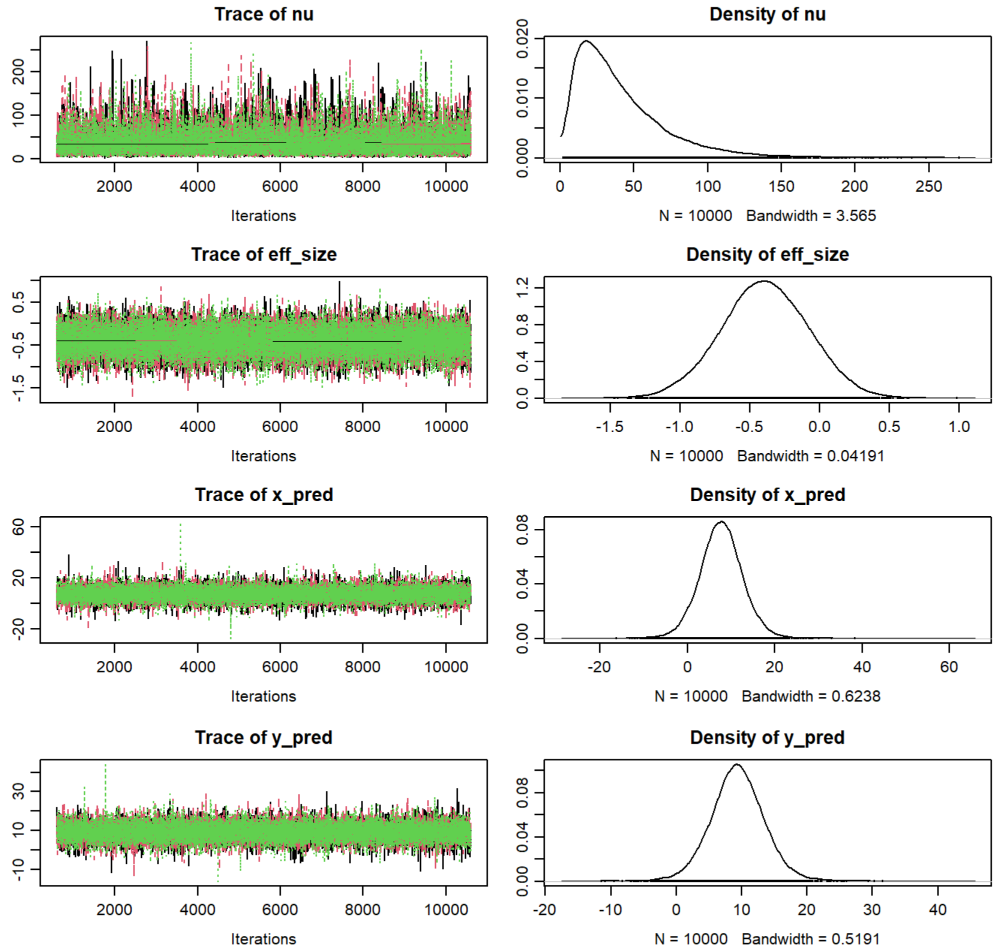


Figure 16: *Bayesian t-test diagnosis plots for the duration data (continued)*

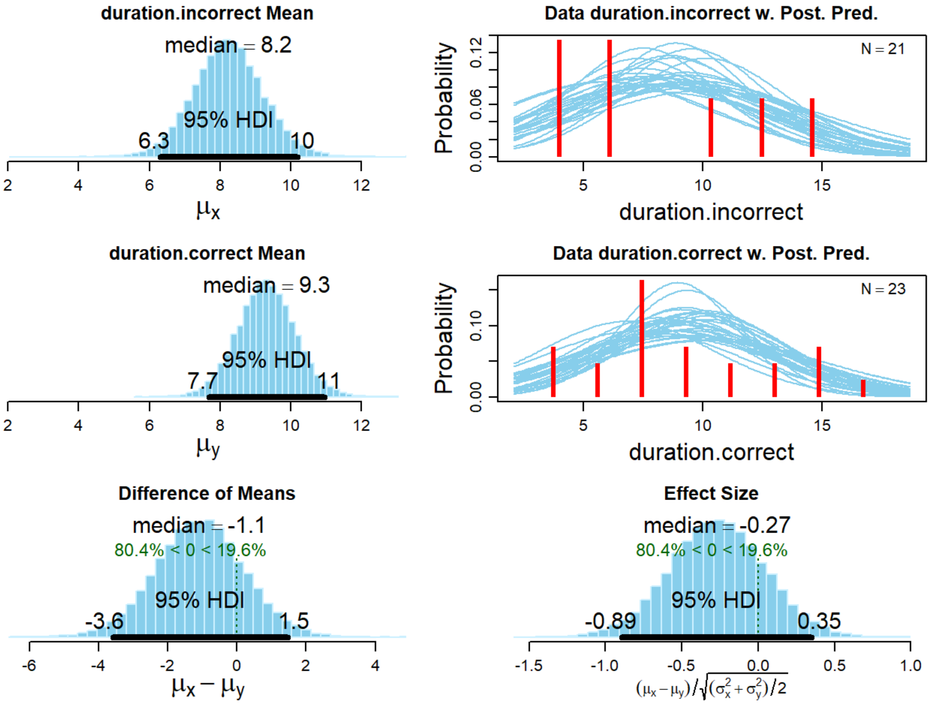


Figure 17: *Bayesian t-test results for the duration data, when not including the excluded groups(groups 44 & 50 due to short duration)*

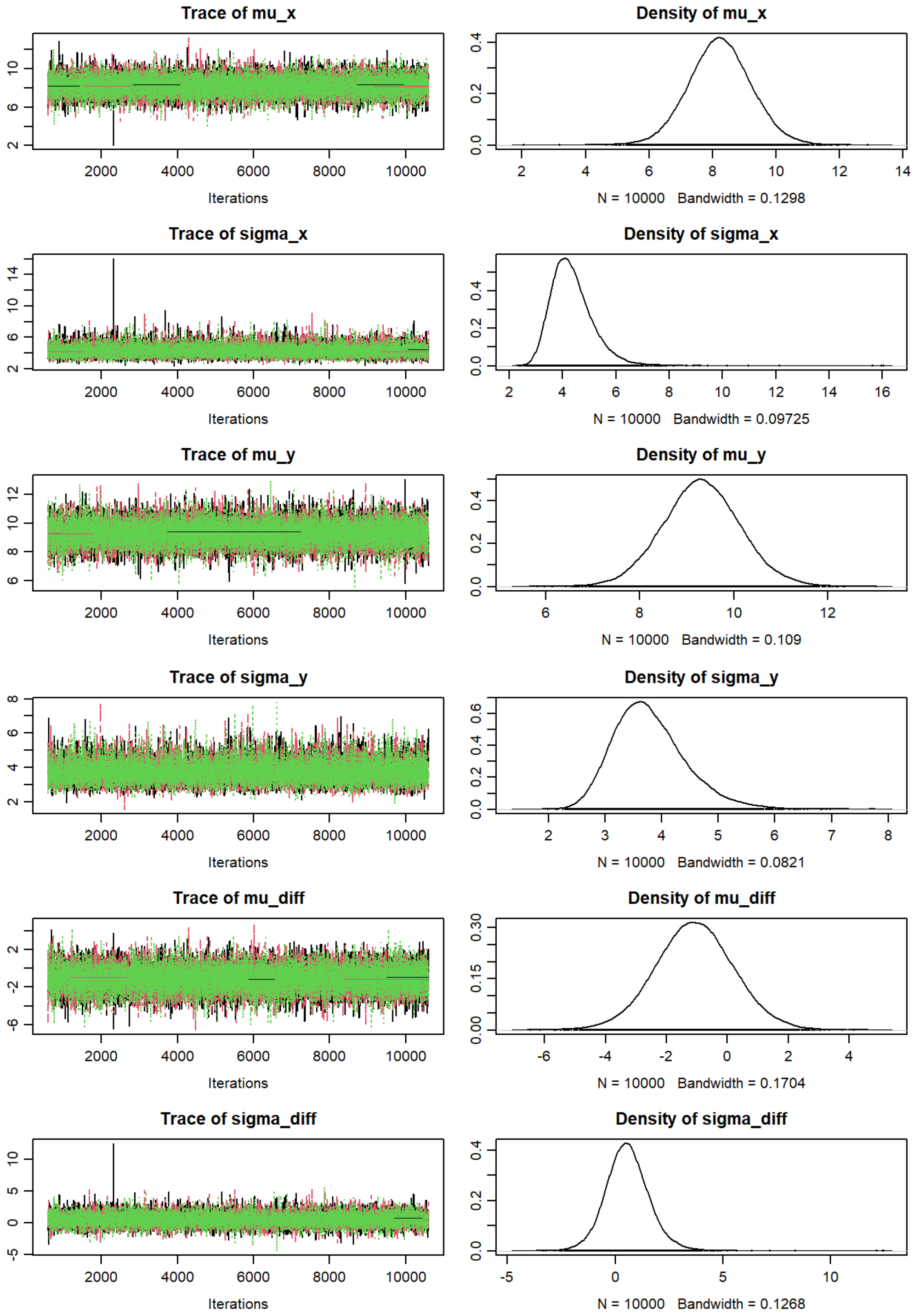


Figure 18: *Bayesian t-test diagnosis plots for the duration data, when not including the excluded groups (groups 44 & 50 due to short duration)*

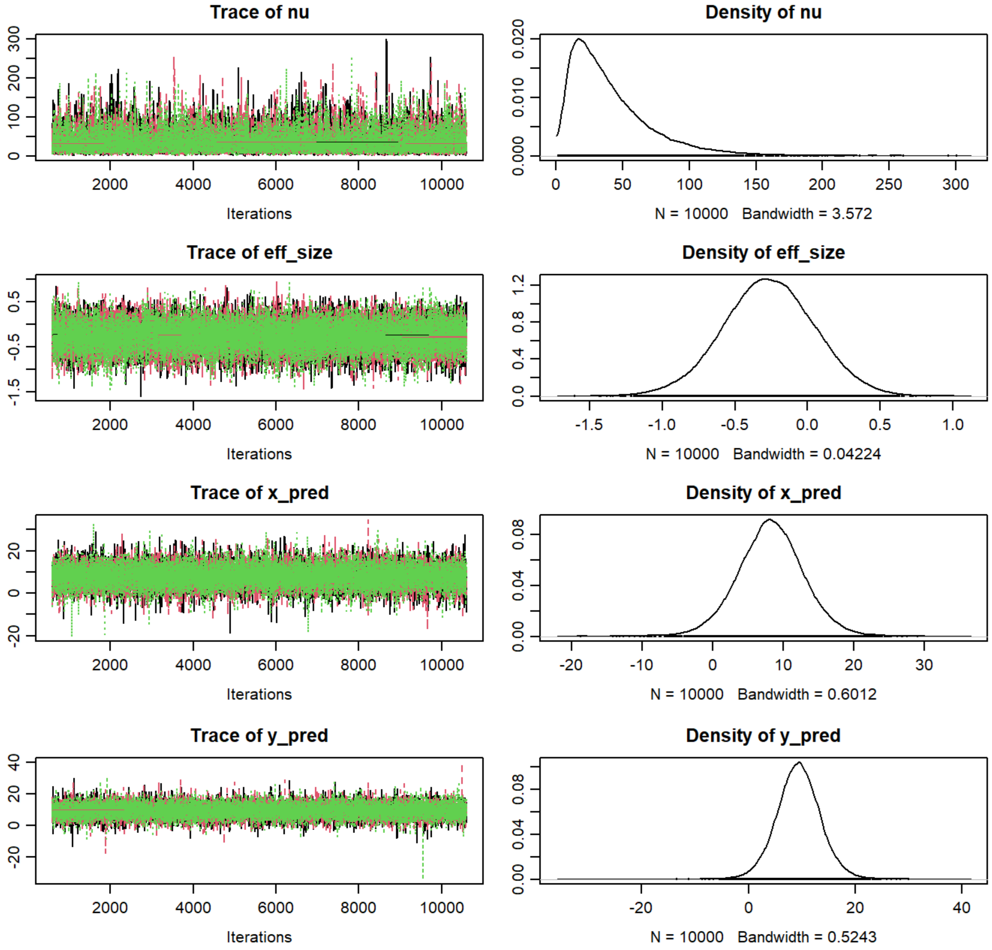


Figure 18: *Bayesian t-test diagnosis plots for the duration data (continued)*

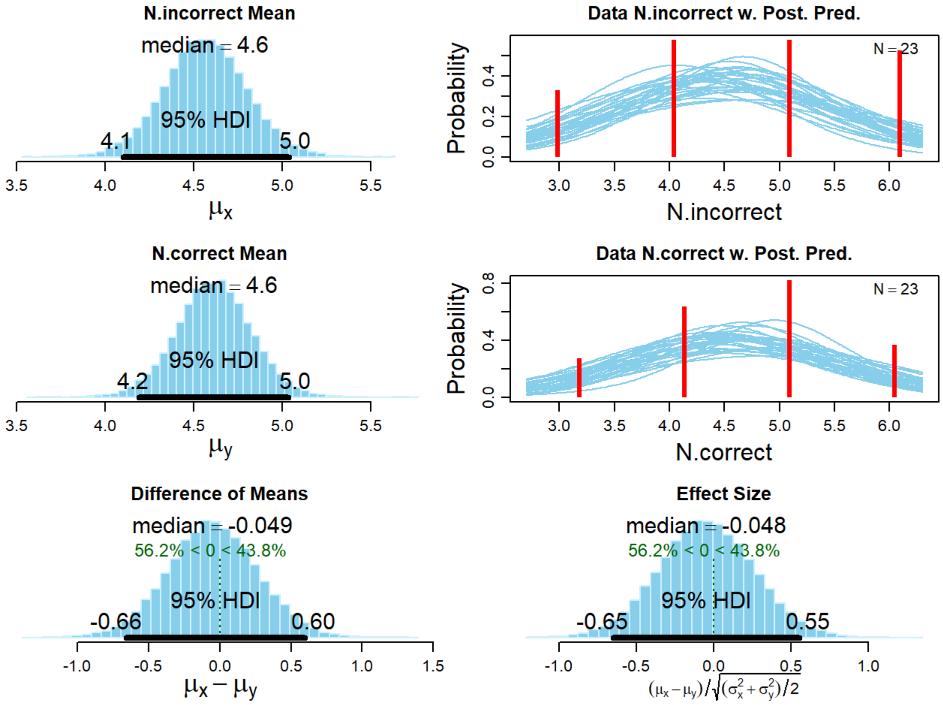


Figure 19: *Bayesian t-test results for the group size data, when including the excluded groups(groups 44 & 50 due to short duration)*

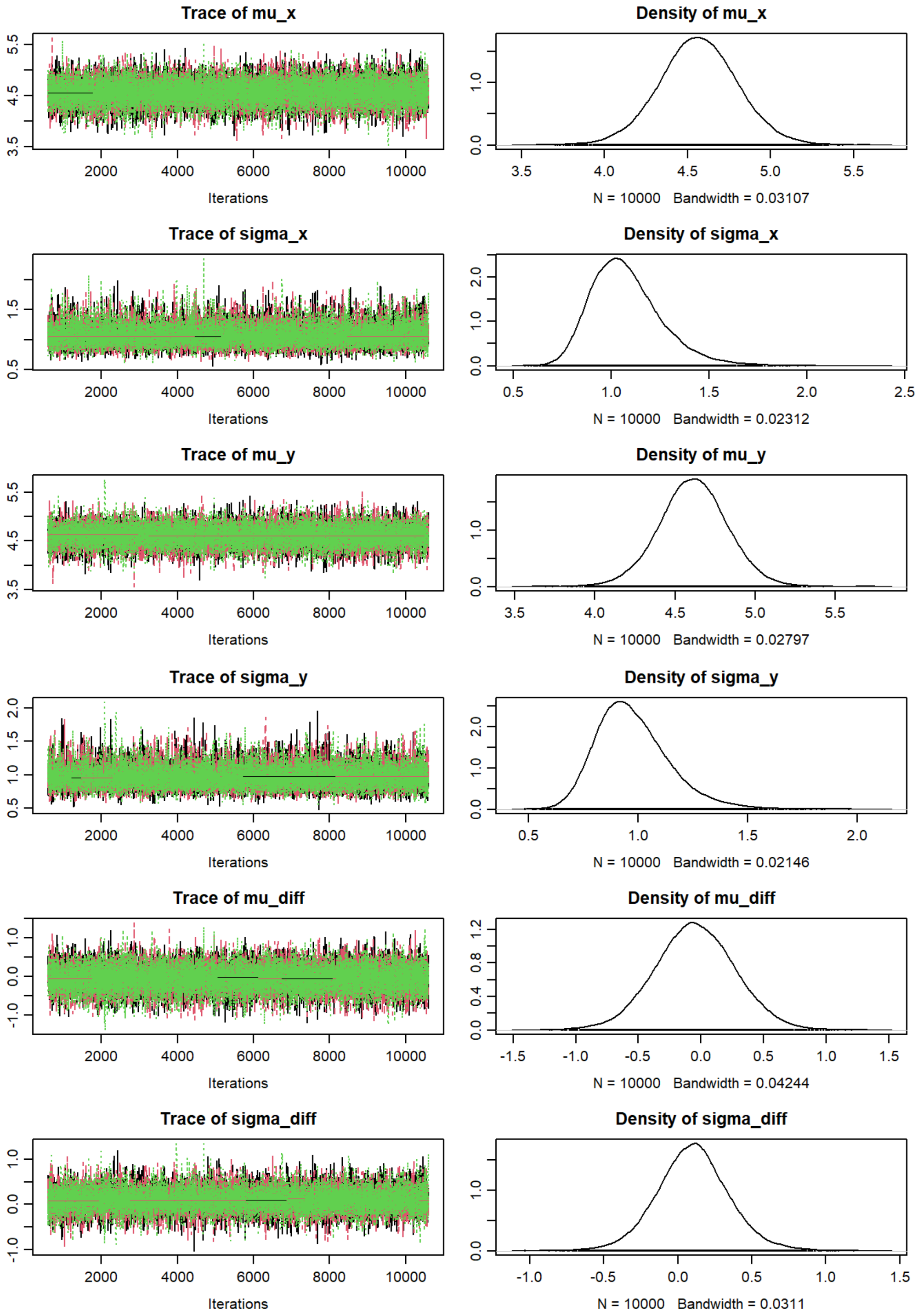


Figure 20: *Bayesian t-test diagnosis plots for the group size data, when including the excluded groups (groups 44 & 50 due to short duration)*

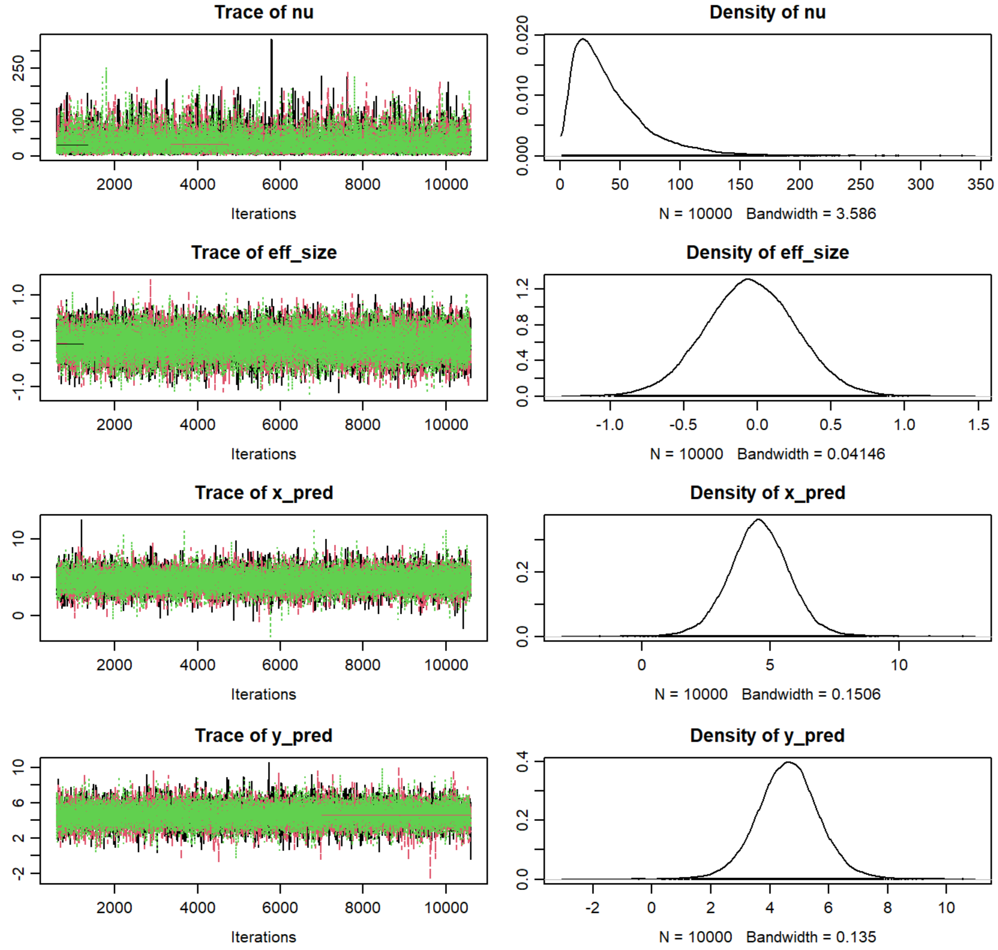


Figure 20: *Bayesian t-test diagnosis plots for the group size data (continued)*

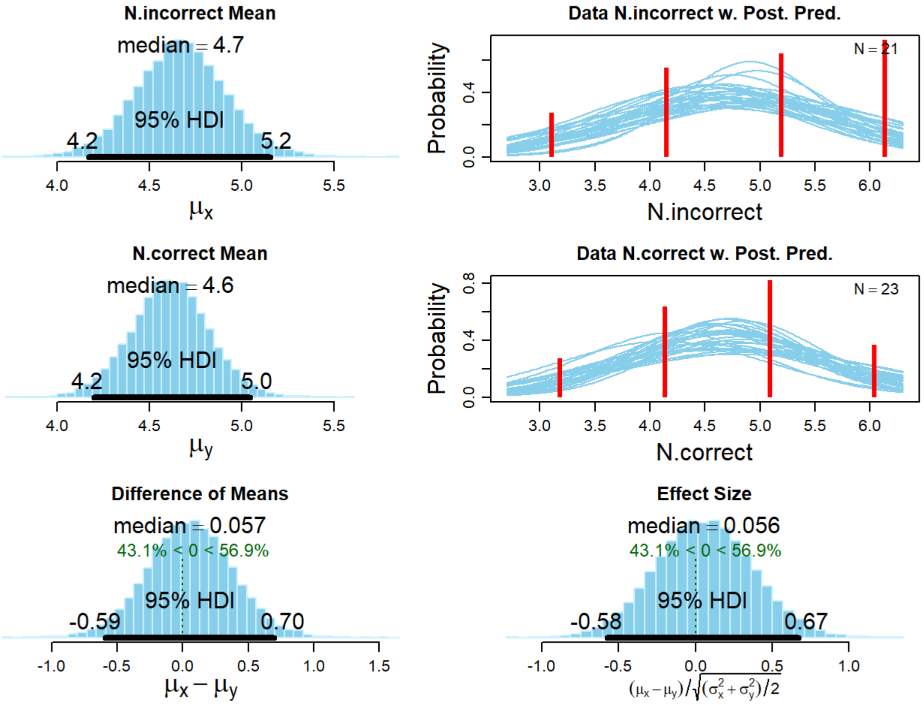


Figure 21: *Bayesian t-test results for the group size data, when not including the excluded groups(groups 44 & 50 due to short duration)*

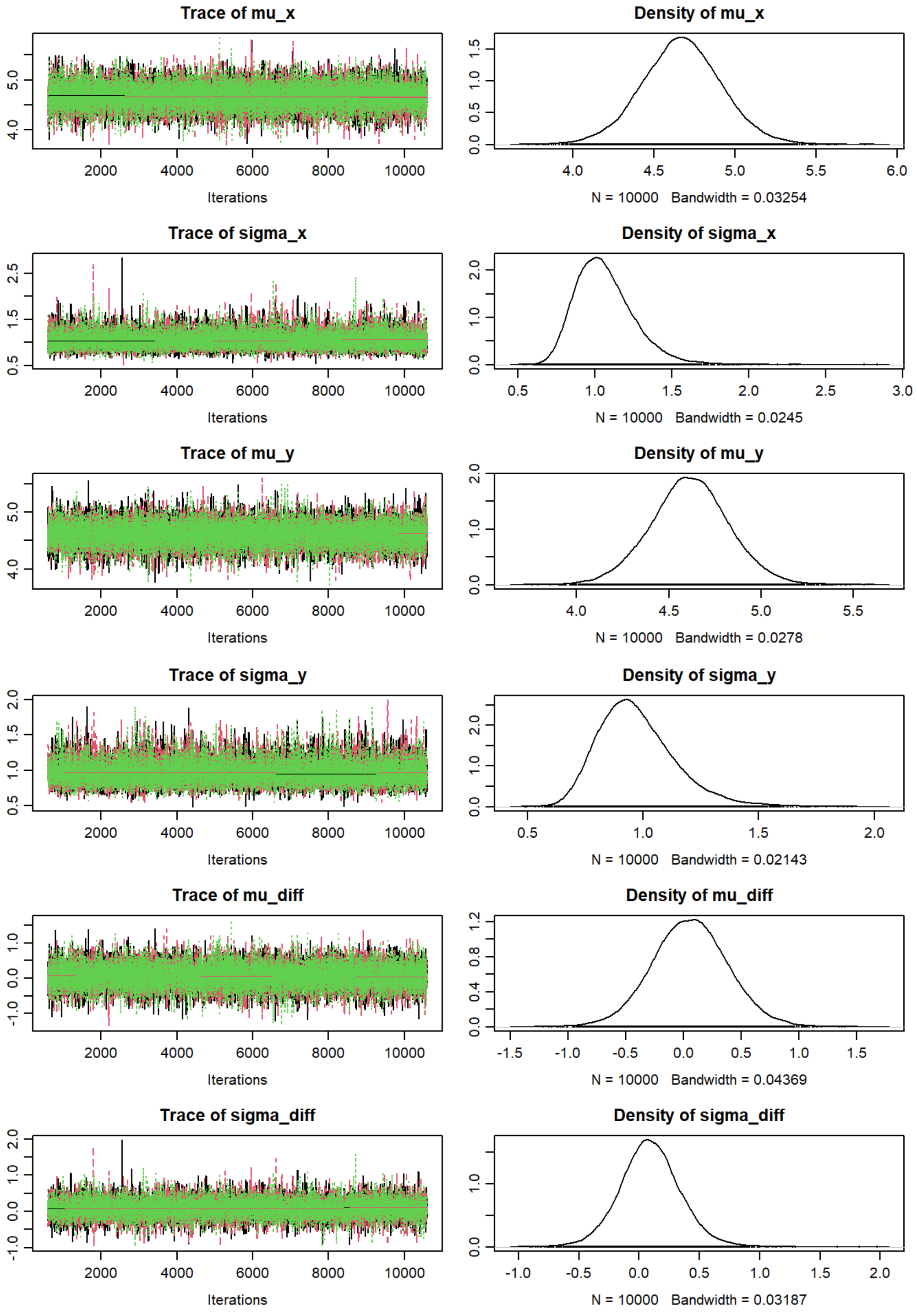


Figure 22: *Bayesian t-test diagnosis plots for the group size data, when not including the excluded groups (groups 44 & 50 due to short duration)*

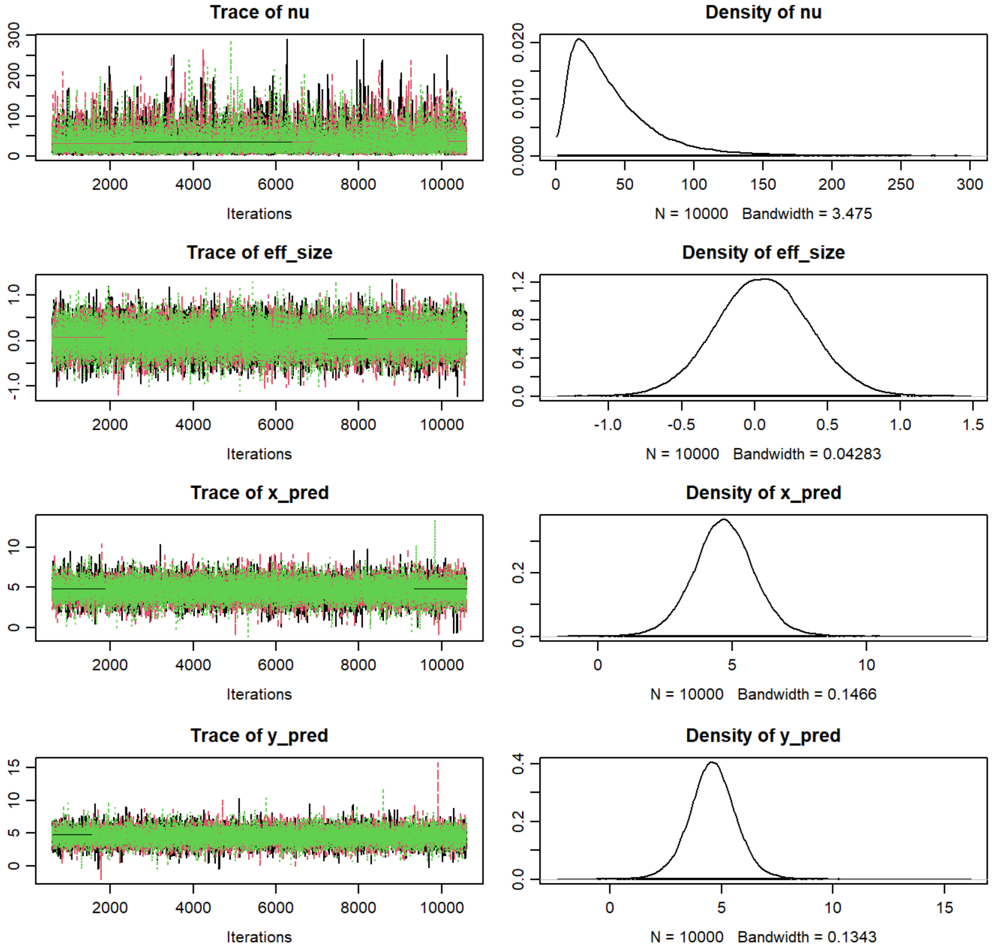


Figure 22: *Bayesian t-test diagnosis plots for the group size data (continued)*
